# Repressor sequestration activates translation of germ granule localized mRNA

**DOI:** 10.1101/2023.10.17.562687

**Authors:** Ruoyu Chen, William Stainier, Jeremy Dufourt, Mounia Lagha, Ruth Lehmann

## Abstract

Biomolecular condensates organize and compartmentalize biochemical processes within cells^1^. Among these, ribonucleoprotein (RNP) granules are characterized as storage depots for translationally repressed mRNA^2–8^. Whether RNP granules can also activate translation and how such translation is achieved remains unclear^9^. This question is particularly relevant for embryonic germ granules, whose activity has been linked to germ cell fate^10–13^. Here, we use single-molecule imaging to show that embryonic germ cell RNP granules in *Drosophila* are the sites of active translation for *nanos* mRNA. Translating *nanos* mRNA is oriented with the 5’end preferentially at the germ granule surface while the 3’UTR is buried within the granule. Untranslated *nanos* mRNAs remain internal within the granule. Quantitative analysis of translational kinetics demonstrates that germ granules activate translation by antagonizing translational repression rather than changing the rate or efficiency of translation. We generated separation-of-function mutations in the disordered linker region of the scaffold protein Oskar that specifically impede *nanos* translation without affecting germ granule morphology or RNA localization. These mutations reveal that *nanos* translation is dependent on the sequestration of translational repressors within germ granules. Together, our findings show that RNP granules regulate localized protein synthesis through compartmentalized relief of translational repression raising the possibility that similar repressor-activator switches control translation in other condensates.

## Introduction

Biomolecular condensates compartmentalize the intracellular environment and biochemical processes to promote efficiency, achieve specificity, and allow regulation at the spatiotemporal levels ^1^. Ribonucleoprotein (RNP) granules are a type of condensate that serves as hubs of post-transcriptional regulation by localizing specific RNAs and RNA binding proteins (RBPs) ^3,4,9^. Most of the well-studied RNP granules, like stress granules ^10^, processing bodies ^2,5^, and neuronal transport granules ^6^, mainly assemble and store translationally repressed mRNA. The assembly of RNP granules can directly cause translational repression ^11–13^. Translation resumes only when the stored mRNAs are released from the RNP granules or the granules undergo disassembly ^7,8,14–16^. Conversely, it has been elusive but curious whether RNP granules can also activate the translation of stored mRNA ^9^. Recently, evidence has emerged that condensation of specific RBP via liquid-liquid phase separation can activate the translation of their target mRNAs ^17^. Several studies have reported that specific cytoplasmic RNP granules may serve as translation factories ^18–21^. However, the role of RNP granules in translational activation remains unclear. Mechanistically it is unknown whether translation occurs on the granule, how the translated mRNA is organized, and whether this involves specific translational regulatory mechanisms acting on the efficiency and rate of translation.

In early *Drosophila* embryos, specialized RNP granules, called germ granules, located in the posterior cytoplasm (also known as germplasm, Fig. 1a) are essential for the formation of primordial germ cells (PGCs) ^22–24^. Several maternally deposited mRNAs (e.g., *nanos*, *gcl,* and *pgc*) crucial for anterior-posterior patterning of the embryo and PGC specification are concentrated in the germplasm. Their translation is restricted to the germplasm even though these mRNAs are also present throughout the entire embryo ^25–31^. It has therefore been proposed that germ granules may serve as specialized compartments for the translation of these mRNAs ^21,28,31^. Thus, Drosophila germ granules serve as a model to gain insight into how RNP granules can control not only the storage but also the activation of translationally silenced RNAs.

**Figure 1.**
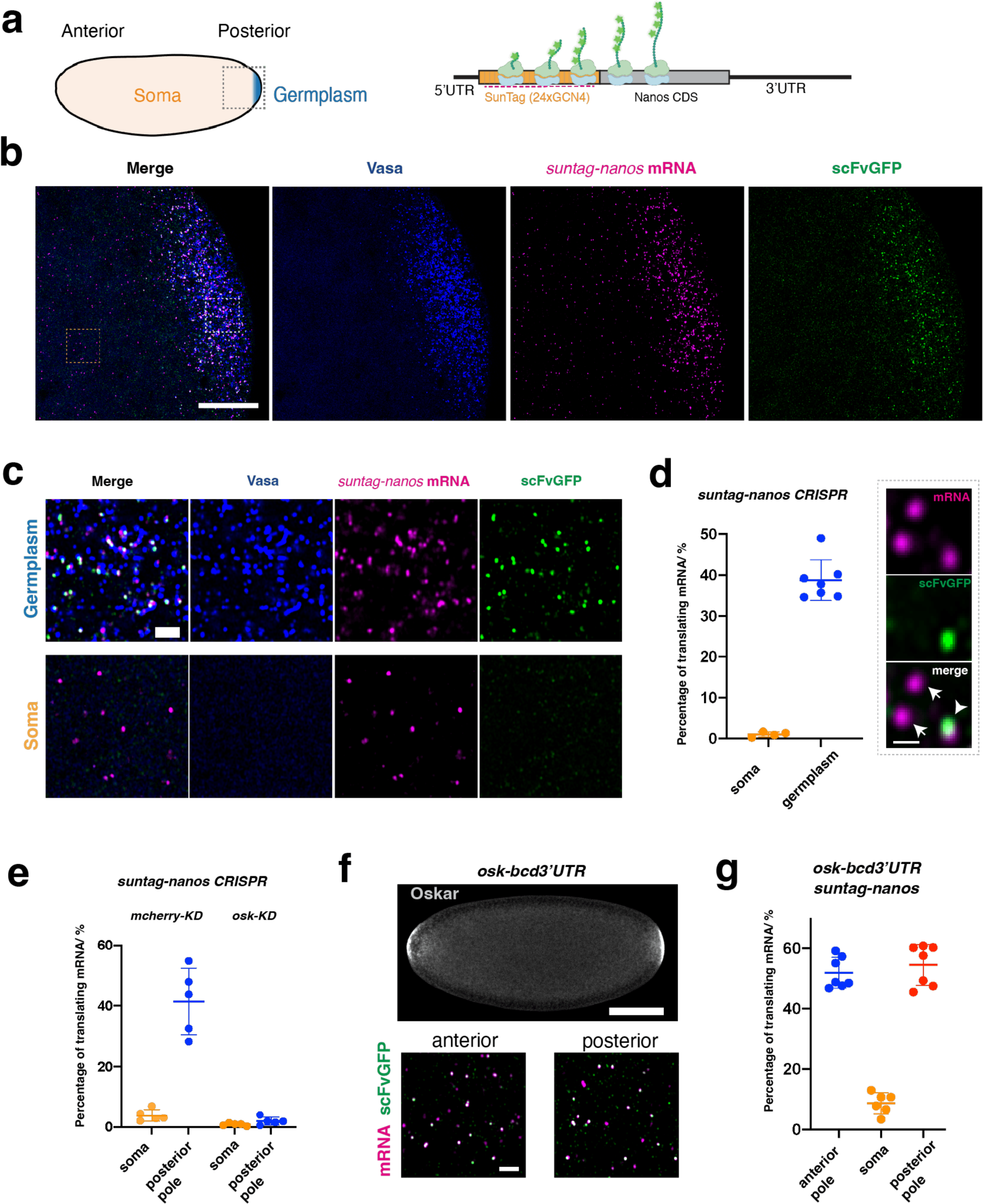
Imaging translation of *nanos* mRNA in Drosophila embryos. **a**, (Left) schematic of a Drosophila embryo. Germplasm (blue) is located at the posterior pole of the embryo. The dashed square represents the region imaged by confocal microscopy and presented in panel **b**. (Right) Schematic of a translating *suntag-nanos* mRNA. A repetitive array of SunTag epitopes is added to the N-terminus of the *nanos* coding sequence. Nascent SunTag peptides are detected by scFv-GFP binding and *suntag* mRNA is detected by smFISH probes (magenta dashed line). **b**, A representative confocal image of the posterior pole of an embryo expressing Vasa-mApple (blue), *suntag-nanos* (mRNA stained by suntag smFISH probes, magenta), and scFv-GFP (green). Outlined regions in germplasm and soma are magnified and presented in panel (C). Scale bar 20 µm. **c**, Magnified images of germplasm and soma show the different translation activities in these two parts of the embryo. Scale bar 2 µm. **d**, (Left) quantification of the percentage of translating mRNA in the soma and the germplasm of embryos. (Right) zoomed confocal images showing examples of a translating mRNA that co-localizes with scFv-GFP signal (arrowhead) and two non-translating mRNA which do not co-localize with scFv-GFP signal (arrows). **e**, Quantification of *suntag-nanos* mRNA translation in the soma and the posterior pole of embryos with *mCherry* (control) knock-down (KD) or *osk* KD. **f**, *Osk-bcd 3’UTR* expression induces germplasm and translation of *suntag-nanos* mRNA at the anterior pole. (Top) Oskar protein is immunostained with anti-Oskar antibody. (Bottom) translation of *suntag-nanos* mRNA in native germplasm at the posterior and induced germplasm at the anterior, which are quantified in **g**. Scale bar 100 µm (F top), 2 µm (F bottom).

Translational repression of *nanos* mRNA in the embryonic soma is mediated by the RNA-binding protein Smaug, which binds to the *nanos* 3’UTR and recruits the translational repressors Cup, an eIF4E-binding protein, and the CCR4-NOT deadenylation complex ^31–34^. De-repression of *nanos* translation at the posterior of the embryo has been attributed to germ granules but the mechanism of de-repression is not well understood ^31,35^. The scaffold protein of germ granules, Oskar (the short isoform), has been proposed to antagonize Smaug’s function ^31,32,36–39^, but the mechanism has not been dissected *in vivo* due to a lack of separation-of-function *oskar* alleles that specifically impede *nanos* translation without affecting germ granule assembly and RNA localization. In this study, we focused on the translational regulation of *nanos* mRNA by germ granules. By direct visualization of *nanos* translation at the single-molecule level using the SunTag technique, we demonstrate that germ granules are the exact sites of *nanos* translation. Taking advantage of the quantitative nature of the SunTag system and a newly generated separation-of-function *oskar* allele that we identified, we dissected the mechanism of translational activation by germ granules *in vivo* at the molecular level.

### *SunTag* system faithfully recapitulates translational regulation of *nanos in vivo*

To investigate whether germ granules are compartments for active translation, we sought to visualize *nanos* translation *in vivo* at the single-molecule level. To this end, we employed the SunTag system, whereby a repetitive array of a GCN4 epitope (SunTag) is appended to the coding sequence of the gene-of-interest and a GFP-fused single chain antibody fragment (scFv-GFP) which binds the GCN4 epitope is co-expressed ^40–44^. By microscopy, the binding of scFv-GFP to the GCN4 epitopes renders nascent peptides emerging from the polysomes as bright GFP foci. Translation of the SunTag can be correlated simultaneously with the corresponding mRNA signal visualized by single-molecule fluorescence *in situ* hybridization (smFISH) (Fig. 1a). Using CRISPR, we knocked a SunTag with 24 copies of the GCN4 epitope into the amino terminus of the endogenous *nanos* coding sequence, referred to as *suntag-nanos* (Extended Data Fig. 1a, see methods and supplementary notes). We utilized a newly developed monomeric msGFP2-fused scFv, which prevents the aggregation of fully synthesized SunTag proteins seen with the original super-folder GFP-fused scFv (Extended Data Fig. 1, b and c) ^45^. Embryos were collected and fixed from female flies carrying *suntag-nanos*, germline-expressing *scFv-GFP,* and *vasa-mApple* as a germ granule marker. The mRNA of *suntag-nanos* was hybridized using smFISH probes against the *suntag* sequence, and the embryos were imaged with confocal microscopy. The mRNA of *suntag-nanos* showed a global distribution within the embryo similar to that of the native *nanos* mRNA: present throughout the embryo while enriched in the germplasm (Fig. 1b). Strikingly, we observed a significant amount of GFP foci in the germplasm, while GFP foci were scarce elsewhere (referred to as soma) (Fig. 1b, c). A zoomed-in view showed that most of the GFP foci were colocalized with smFISH foci, representing individual translation sites (Fig. 1c, d). Injection of puromycin, a translation inhibitor that disassembles polysomes, abolished the GFP foci, validating that the GFP foci represented actively translating polysomes (Extended Data Fig. 1d). We used FISH-QUANT to locate individual RNA foci and GFP foci and determine whether an RNA molecule colocalized with a GFP focus, thus being translated ^46^. We detected 30%∼50% *suntag-nanos* mRNA being translated in the germplasm (Fig. 1c, d), whereas in the soma, the percentage was lower than 2% on average (Fig. 1c, d). Detecting SunTag in stage 1 embryos using anti-GCN4 immunostaining instead of scFv-GFP provided comparable results (Extended Data Fig. 2). The SunTag immunostaining signal resembled the spatial pattern of Nanos protein immunostaining in a wildtype (WT) embryo (Extended Data Fig. 2c). At stage 5 when PGCs are fully cellularized, there was a strong reduction of bright scFv-GFP foci within PGCs, seemingly suggesting that translation was repressed (Extended Data Fig. 2d, e). Anti-GCN4 immunostaining, however, confirmed that *suntag-nanos* translation was active after PGC formation and the reduction of GFP foci was due to the limitation and depletion of scFv-GFP within PGCs (Extended Data Fig. 2b, d, and e) ^45^. Thus, active *suntag-nanos* translation was maintained throughout PGC formation (Extended Data Fig. 2b), consistent with the accumulation of Nanos protein in PGCs seen in previous studies ^30,47^. Together, *suntag-nanos* mRNA exhibited a localized translation pattern in the germplasm that is consistent with the translation pattern described for the native *nanos* mRNA.

Translation of *nanos* mRNA in the germplasm is dependent on the assembly of germ granules and can occur at the anterior pole of an embryo if germ granules are ectopically formed there ^35,47^. In agreement, translation of *suntag-nanos* at the posterior pole was abolished when germ granule assembly was perturbed by knocking down maternal *oskar* expression (Fig. 1e, and Extended Data Fig. 3a). We induced germ granule assembly at the embryo’s anterior pole by expressing Oskar protein via transgenic *osk-bcd3’UTR* ^47^. The mRNA of *suntag-nanos* localized to the anterior pole similar to the native *nanos* and was translated at level comparable to native germplasm at the posterior (Fig. 1f, g, and Extended Data Fig. 3b), validating the necessity and sufficiency of germ granules in activating *nanos* translation.

*Nanos* mRNA is synthesized during oogenesis and becomes localized to the germplasm in developing oocytes ^30^. To further validate the translational regulation on *suntag-nanos*, we assessed its translation during oogenesis (Extended Data Fig. 4a). Imaging translation in the germplasm of mature oocytes showed a significantly lower translation rate of *suntag*-*nanos* than in the embryonic germplasm (Extended Data Fig. 4b, c), which is expected due to the widespread translational dormancy of mature oocytes followed by translational activation after egg activation ^30,48,49^. Furthermore, recapitulating egg activation *in vitro* by immersing mature oocytes in a hypotonic buffer was sufficient to restore active translation in the germplasm (Extended Data Fig. 4d, e, and supplementary movie 1) ^49^. In contrast, *suntag-nanos* translation in nurse cells is constitutively active (Extended Data Fig. 4b, c), consistent with the abundance of Nanos protein in nurse cells observed previously ^30,50^. Thus, our results show that *suntag-nanos* faithfully recapitulates the spatiotemporal pattern of translation described by prior studies for the native *nanos* mRNA. This establishes *suntag-nanos* as a reliable tool to study translational control in germ granules.

### Translating mRNAs are positioned in 5’-3’ orientation on germ granules

Using *suntag-nanos* as a single-molecule visual translation reporter *in vivo*, we mapped the distribution of *nanos* mRNA translation relative to germ granules. To this end, we established an image analysis pipeline to measure the distance between individual mRNA or GFP foci and their closest granule border. We first defined the border of individual germ granules by segmenting the Vasa signal with Ilastik, a machine-learning-based image analysis program (Extended Data Fig. 5a) ^51^. The coordinates of individual mRNA and GFP foci were determined and extracted using FISH-Quant and their relative distance to the closest granule border was mapped (Extended Data Fig. 5b, see methods). Consistent with the fact that *nanos* mRNA localizes to germ granules and that low-abundance mRNAs tend to reside at the border of granules ^52,53^, we found that *suntag-nanos* mRNA (detected by *suntag* smFISH) was enriched around the granule border. As controls, we performed the same analysis on simulated random spots, randomized *suntag-nanos* mRNA foci by rotating the image of the smFISH channel, or smFISH foci of *osk* mRNA, which do not localize to germ granules ^52,54^. Foci in all three control images exhibited a similar distribution that was not enriched around the border (Extended Data Fig. 5, c to g). These results demonstrate that the distribution of *suntag-nanos* mRNA is non-random and centered around the granule border, confirming previous single-molecule studies ^52,53,55^.

Next, we analyzed the distribution of GFP foci to infer the position of polysomes relative to the granule border. GFP foci (i.e., polysomes) were enriched around the granule border, which overlapped significantly with the distribution of the *suntag* smFISH spots, consistent with their close physical association (Fig 2a, b). This result demonstrates that *nanos* translation occurs close to the border of germ granules. To corroborate this finding, we immunostained a ribosomal protein RPS6 as a proxy for ribosomes (Extended Data Fig. 6). Within germplasm, RPS6 staining exhibited an enrichment within and around germ granules (Extended Data Fig. 6), supporting the model that germ granules are the translation ‘hotspots’ within germplasm ^56^. When *suntag-nanos* mRNA was alternatively detected with smFISH probes targeting the *nanos* 3’UTR, we noticed a shift between the smFISH and GFP distributions, reflecting the separation between the coding sequence (CDS) of mRNA and 3’UTR (Fig. 2c). Interestingly, relative to the GFP spots, *nanos* 3’UTR smFISH spots were skewed toward the inside of the granule (Fig. 2d), suggesting that the 3’UTR of translating *suntag-nanos* mRNA is preferentially embedded inside germ granules. This *in vivo* observation is consistent with the *nanos* 3’UTR being necessary and sufficient for mRNA localization to germ granules and the interaction between the *nanos* 3’UTR and RNA binding proteins (Oskar and Aubergine) in germ granules ^28,37,39,57–59^. Taken together, our analysis revealed a specific conformation adopted by germ granule-localized, translating *nanos* mRNA: the CDS and associated polysome of translating mRNA are oriented toward or exposed on the surface of granules, while the 3’UTR is anchored internally (Fig. 2i).

**Figure 2.**
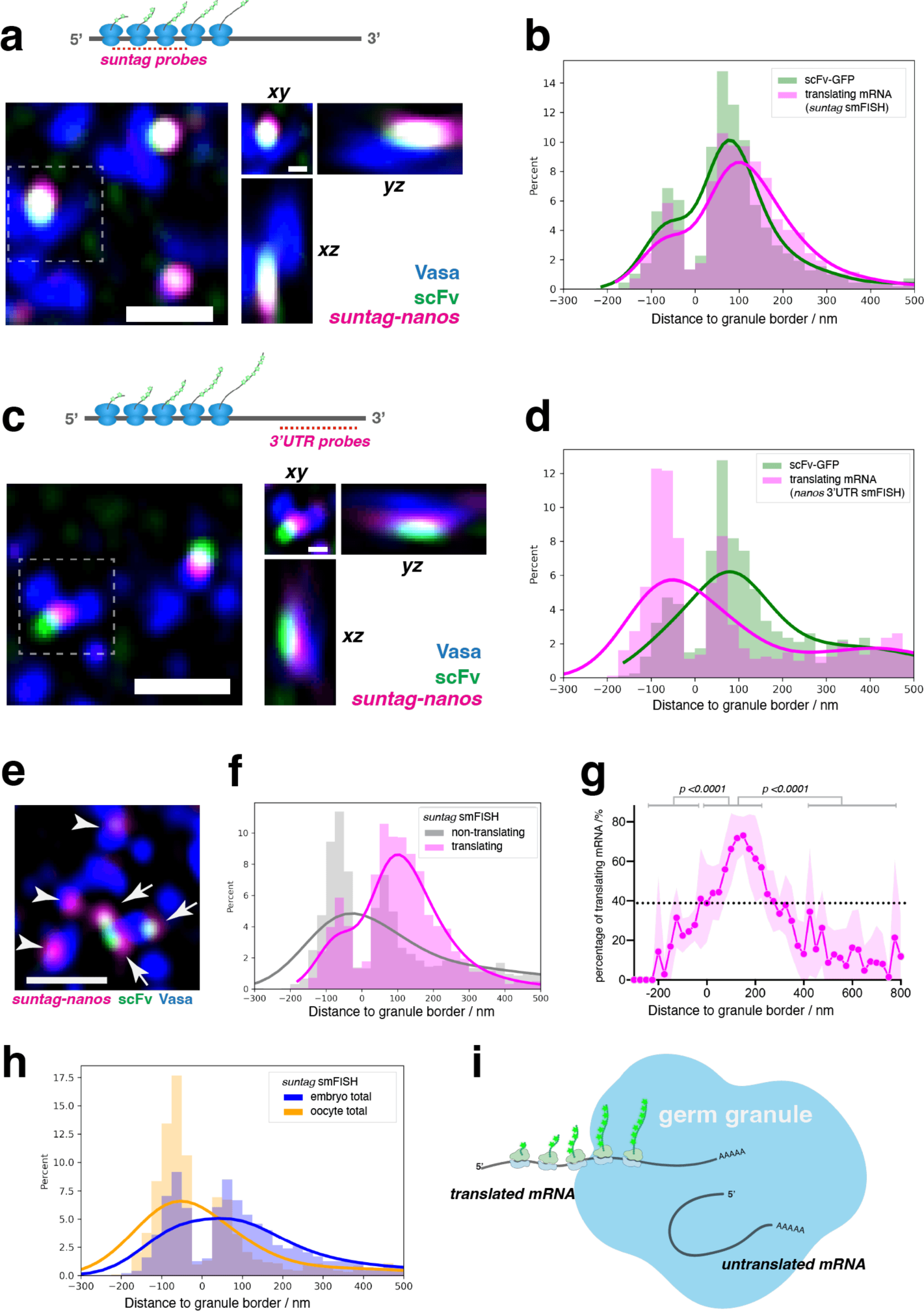
Spatial distribution of the polysome and orientation of translating mRNA. **a-d**, Orientation of translating mRNA. Translating *suntag-nanos* mRNAs in embryos from Vasa-mApple/+; *suntag-nanos*, scFv-GFP/*Df(nanos)* flies (see methods) are detected with smFISH against *suntag* (**a** and **b**) and *nanos* 3’UTR (**c** and **d**). Example germplasm images are shown in **a** and **c**. Scale bar 1 µm. The orthogonal views of outline regions are shown on the right. Scale bar 0.3 µm. Blue, Vasa; magenta, mRNA smFISH; green, scFv-GFP. The distributions of scFv-GFP and smFISH foci were mapped and plotted in relative frequency histograms overlaid with kernel density estimate (KDE) in **b** and **d**. The x-axis refers to the distance of foci centroids to the border of the closest granule; the zero marks granule border; a negative value denotes being inside a granule and positive denotes outside. In total, 12684 smFISH foci and 12733 scFv-GFP foci from 7 images were mapped in **b**. 5663 smFISH foci and 5649 scFv-GFP foci from 3 images were mapped in **d**. **e**-**g**, Polysomes distribute preferentially on the surface of germ granules. **e**, Example image showing the distribution of translating (arrows) and non-translating (arrowheads) mRNA stained by *suntag* smFISH probes. Blue, Vasa; magenta, *suntag* mRNA; green, scFv-GFP. Scale bar 1 µm. **f**, Relative frequency histogram with KDE of translating and non-translating mRNA distribution in germplasm. 12684 translating and 19712 non-translating foci from 7 images were plotted. **g**, The average translating fraction in each bin of the x-axis from 7 images was calculated and plotted. The average translating fraction in entire germplasm is indicated as the dashed line. The translating fractions on the granule surface (0 ≤ x ≤ 200 nm) were compared with the ones within granules (x < 0) or ones not localized to granules (x > 400 nm) using Welch’s t-test. **h**, Distribution of total *suntag-nanos* mRNA stained by *suntag* probes in stage 1 embryos and stage 14 oocytes. 6468 foci from 4 oocyte images and 32473 foci from 7 embryo images were mapped. **i**, A model of the predicted orientation and distribution of translating and non-translating mRNAs in germ granules.

Next, we asked how the translational status of mRNA affected its distribution in germ granules by comparing the distributions of translating versus non-translating *suntag* smFISH foci (Fig. 2e). We noticed that *suntag* smFISH foci of non-translating mRNAs distributed more toward the inside of granules, a pattern similar to that of the distribution of the 3’UTR of translating mRNA (Fig. 2f), suggesting that the 5’UTR and CDS of mRNAs appeared to reside inside granules when the mRNA was not being translated (Fig. 2i). In contrast, translating mRNAs tended to have the CDS localized to the surface of germ granules (Fig. 2f, g). Consistent with this model, in oocyte germplasm where *suntag-nanos* translation was largely repressed, *suntag* smFISH foci distribution showed a global inward shift compared to embryonic germplasm (Fig. 2h, and Extended Data Fig. 4c). To test the causality between translation and exposure of the CDS, we used puromycin to disassemble polysomes in the germplasm. We did not detect any inward shift of *suntag* smFISH signal distribution after puromycin treatment, suggesting translation or polysomes *per se* were not required to maintain the outward orientation of mRNA CDS (Extended Data Fig. 7). Instead, a yet unknown mechanism may drive and sustain the orientation of the CDS toward the granule margin during granule-dependent translation.

### Germ granules de-repress translation without affecting initiation and elongation rates

After establishing that germ granules were the sites of *nanos* translation, we investigated the mechanism of translational activation in germ granules. The translational repression of *nanos* in the soma is mediated by the translational repressor Smaug which binds to the Smaug Response Element (SRE) in the *nanos* 3’UTR ^32,60,61^. To explore Smaug-mediated translational regulation, we generated transgenic flies with *UAS-driven suntag-nanos* constructs that varied in their 3’UTRs. Driven by a maternal Gal4 activator, the respective RNAs either carried a wildtype *nanos* 3’UTR (*suntag-nanos-WT*), which showed germplasm-restricted translation, the same pattern as the CRISPR-generated *suntag-nanos;* a 3’UTR with a mutated SRE (*suntag-nanos-SREmut*) that directed RNA localization to the germplasm but lacked binding sites for the Smaug repressor and exhibited significantly elevated translation in the soma; and a *tubulin* 3’UTR that was evenly distributed throughout the embryo did not bind Smaug, and supported constitutive translation in embryos (*suntag-nanos-tub3’UTR*) (Fig. 3a, b, and Extended Data Fig. 8). Consistent with previous studies, these *suntag-nanos* constructs confirmed the requirement of the *nanos* 3’UTR for RNA enrichment in granules and the role of the SRE sequence for translational repression in the soma ^32,60,61^.

**Figure 3.**
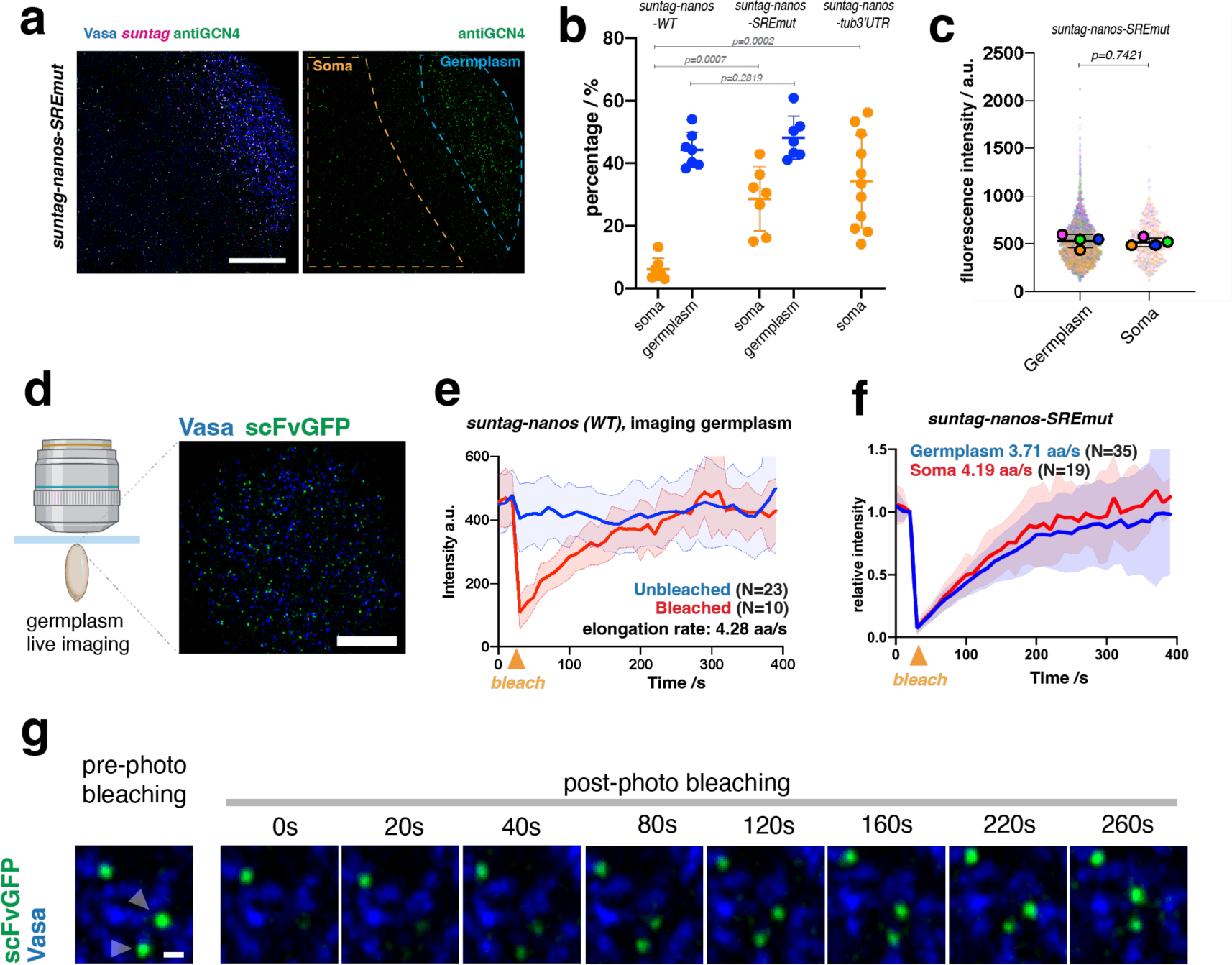
Kinetics of *suntag-nanos* translation. **a**, Example image of the posterior of an embryo expressing Vasa-mApple (blue), *nanos-suntag*-SREmut (mRNA stained by *suntag* probes, magenta). SunTag is stained with anti-GCN4 (green). An image of the anti-GCN4 channel is shown on the right with germplasm and soma outlined, showing that translation is prevalent in both soma and germplasm. Scale bar 20 µm. **b**, Quantification of the translating fractions in embryos from flies expressing transgenic *suntag-nanos-WT*, *suntag-nanos-SREmut*, or *suntag-nanos-tub3’UTR*. Stage-2 embryos were used for quantification. Pairwise statistical comparisons were conducted using Welch’s t-test. **c**, The intensities of polysomes (anti-GCN4 staining) in soma and germplasm of embryos from flies expressing *UAS-suntag-nanos-SREmut*. Quantification results from four embryos were plotted in a super-plot. Individual dots represent the intensities of individual polysomes, each color-coded by embryos. Each colored circle represents the mean intensity of each embryo. The black lines and error bars indicate the mean and standard deviation (SD) of the four embryos. Statistical comparison was performed on the mean intensities of individual embryos using Welch’s t-test. **d**, (Left) Schematic of the live imaging setup. The posterior pole of the live embryo is stuck onto a coverslip and imaged by an upright microscope. (Right) a representative image of germplasm during live imaging. Blue, Vasa; green, scFv-GFP. Scale bar 10 µm. **e**, The intensities of polysomes over time during live imaging *suntag-nanos* mRNA translation with (red curve, average of 10 curves) or without (blue curve, average of 23 curves) photo-bleaching when Time= 30s. The elongation rate calculated from the plot is indicated. **f**, Polysome intensities of *suntag-nanos-SREmut* mRNA in germplasm (blue curve, average of 35 curves) and soma (red curve, average of 19 curves) over time with photo-bleaching when Time= 30s. The elongation rates in germplasm and soma calculated from the plot are indicated. **g**, Representative time-lapse image of fluorescence recovery after photobleaching (FRAP) of two translation sites (arrowheads). Blue, Vasa; green, scFv-GFP. Scale bar 500 nm.

We used these constructs to analyze translational dynamics quantitatively at the single-molecule level. This revealed that, in germ granules, a similar fraction of *suntag-nanos-SREmut* mRNA was translated compared to *suntag-nanos-WT* (Fig. 3b). The fact that local translation was not significantly increased in the Smaug binding site mutant implies that SRE sequences do not mediate repressor activity in the germplasm ^31,32^. However, the fraction of translating *suntag-nanos-SREmut* mRNA was higher in the germplasm than in the soma (Fig. 3b). One possibility is that, in addition to Smaug, *nanos* translation is suppressed by other translational repressors ^62^, which are not mediated by SRE sequences in the 3’UTR but also counteracted by germ granules. Alternatively, and not mutually exclusive, germ granules may also actively promote translation in addition to de-repression. In line with this hypothesis, we observed an enrichment of eIF4G and PABP with germ granules (Extended Data Fig. 9). Moreover, it has been shown that eIF4A and eIF3 are recruited by the germ granule components Tudor and Aubergine ^21,63^. Thus, in addition to a dominant de-repression mechanism needed to overcome Smaug, select translation factors recruited by germ granules may facilitate mRNA translation. Together these results demonstrate a primary role for germ granules in protecting localized mRNA from translational repression, thereby allowing translation to occur.

Our results show that localized *nanos* mRNA was specifically translated on germ granules, and that this was achieved primarily in germ granules by preventing translational repression. Next, we asked whether germ granules specifically modulate the kinetics of translation ^40,64^. For example, germ granules may boost *nanos* translation by increasing translational initiation or elongation rate, apart from protecting *nanos* from translational repression by Smaug. We utilized the *suntag-nanos-SREmut* transgene to directly compare the translation kinetics of unlocalized mRNA in the soma to that of localized mRNA in granules within the same embryo. We measured the intensity of individual polysomes (SunTag staining) and found that the average intensities did not differ significantly between the germplasm and the soma (Fig. 3c). As polysome intensity correlates with the number of ribosomes loaded onto *suntag-nanos* mRNA, which is determined by the steady-state translational initiation rate ^64,65^, this result indicated that translating *suntag-nanos* mRNA might have similar ribosome occupancy and initiation rate between germplasm and soma. We utilized the intensity of fully synthesized SunTag-Nanos peptide (dim SunTag foci without colocalized mRNA) to estimate the number of ribosomes on a translating mRNA (Extended Data Fig. 10a, b, also see methods) ^40^. Translating *suntag-nanos* mRNA carried about one ribosome per 300 nucleotides in the coding sequence (Extended Data Fig. 10c), which is comparable to reported measurements carried out in tissue culture systems and *Drosophila* embryos ^40–42,65,66^.

To measure the elongation rate of translation in the soma and germplasm, we utilized fluorescence recovery after photo-bleaching (FRAP) of translation foci in live embryos (Fig. 3, d to g) ^64^. We tracked individual translation foci in live embryos for over five minutes (Fig. 3e, supplementary movie 2). Most tracked foci maintained their intensity over the live imaging process, indicating a steady state with constant translational initiation and elongation. After photo-bleaching individual foci, GFP fluorescence recovered over time and plateaued at the initial intensity (Fig. 3, e to g, and supplemental movie 3). It has been reported that the binding of scFv-GFP to SunTag epitopes is stable, with a binding half-life of 5 to 10 min ^67^. The full recovery of translation foci took around 4 min, indicating that the synthesis of new SunTag peptides, instead of exchange of scFv-GFP, led to the fluorescence recovery (Fig. 3e, f). We found that the FRAP curves of GFP spots in the soma and the germplasm closely matched (Fig. 3f, and supplemental movie 3 and 4), suggesting similar elongation rates. We used a mathematical model to fit the FRAP data and calculate the elongation rate (Extended Data Fig. 10d, e, also see methods) ^41,44,64^, which yielded 3.71 aa/s and 4.19 aa/s in the soma and the germplasm respectively. These values are similar to the eukaryotic translation elongation rates calculated from ribosome profiling experiments and SunTag imaging in tissue culture ^40,41,68,69^. Together, these results suggest that germ granules do not increase the steady-state initiation and elongation rates of translation. Instead, our results are consistent with the conclusion that germ granules allow the translation of *nanos* mRNA mainly by counteracting translational repression by Smaug.

### Oskar de-represses *nanos* translation by regulating the localization of translational repressors

It has been unclear how germ granules protect *nanos* mRNA from the repression by Smaug, which binds to the SRE sequences within the *nanos* 3’UTR ^32,61^. As we observed that translating *suntag-nanos* mRNA embedded the 3’UTR inside germ granules, germ granules may create a space excluding Smaug and consequently protect *nanos* 3’UTR from Smaug binding and repression (Fig. 2i). By imaging transgenic Smaug-GFP embryos, we observed that Smaug was present throughout the embryos, forming heterogeneous clusters in the soma (Extended Data Fig. 11a). In the germplasm and PGCs, however, we unexpectedly found that Smaug was enriched within the germ granules, refuting the exclusion model (Fig. 4d, and Extended Data Fig. 11a). It has been established that Smaug represses translation by recruiting the eIF4E-binding protein Cup and the CCR4-NOT deadenylation complex to inhibit translational initiation and assembling a stable repressive RNP complex with the P-bodies protein ME31B (DDX6 homolog) ^34,36,62,70^. We examined the distribution of these Smaug co-factors (Cup, CCR4, NOT3 and ME31B) in germplasm and found that, unlike Smaug, these were not enriched within germ granules (Extended Data Fig. 11b and Extended Data Fig. 12), indicating that the germ granule-localized Smaug appeared unable to recruit the necessary downstream effectors needed for translational repression. Specifically, we found that ME31B was localized on the surface of germ granules after PGC formation, while in the soma ME31B forms distinct micron-size granules (Extended Data Fig. 11b). Thus, germ granule-localized Smaug may not be conducting its role as a translational repressor.

**Figure 4.**
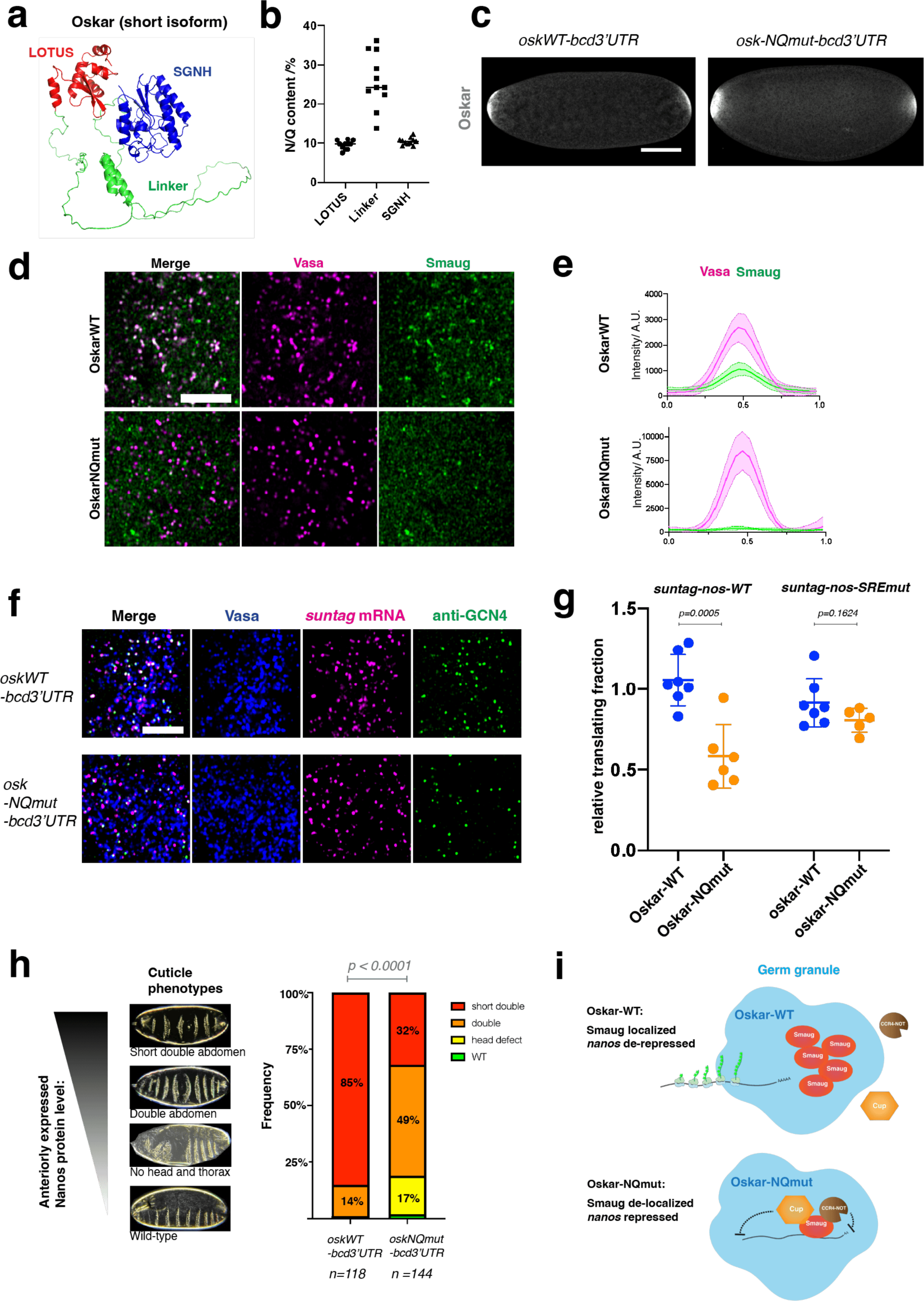
Oskar linker region controls Smaug localization and *nanos* translation. **a**, AlphaFold structure model of short Oskar protein, with LOTUS domain in red, SGNH-like domain in blue, and linker region in green. **b**, Percentage of glutamine (Q) and asparagine (N) in three regions of short Oskar proteins from eleven *Drosophila* species. Each dot represents an Oskar of a particular *Drosophila* species. **c**, *Oskar-NQmut-bcd3’UTR* induces anterior germplasm similarly to *Oskar-WT-bcd3’UTR*. Oskar-WT/NQmut proteins are immunostained with anti-Oskar antibody. Scale bar 100 µm. **d**, Distribution of Smaug in germplasm. Images of induced germplasm by Oskar-WT or Oskar-NQmut at the anterior pole. Germ granules are labeled by Vasa-mApple (magenta). Smaug is visualized with Smaug-GFP (green). Scale bar 5 µm. **e**, Intensity profiles of Vasa-mApple (magenta) and Smaug-GFP (green) along the lines across the germ granules induced by Oskar-WT or Oskar-NQmut. Intensity profiles of 20 germ granules were combined for each genotype, where the curves represent the mean value and the color-filled areas cover the standard deviation. **f**, Representative images showing the translation of *suntag-nanos* mRNA in germplasm induced by Oskar-WT or Oskar-NQmut. Blue, Vasa; magenta, suntag smFISH; green, anti-GCN4. Scale bar 5 µm. **g**, The fraction of *suntag-nanos-WT* or *suntag-nanos-SREmut* mRNA being translated in anterior germplasm induced by Oskar-WT or Oskar-NQmut. Each dot represents the normalized measurement of an embryo where the translating fraction in the anterior germplasm is divided by the translating fraction in the native germplasm at the posterior. Statistical comparisons between Oskar-WT and NQmut were performed by t-test, and p-values are indicated. **h**, The cuticle phenotypes generated by *Oskar-WT/NQmut-bcd3’UTR*. The cuticle images show a range of cuticle phenotypes corresponding to different levels of anteriorly-expressed Nanos protein. The bar graph shows the frequency of each cuticle phenotype caused by *Oskar-WT/NQmut-bcd3’UTR* expression. Statistical comparison was performed using Chi-square test. **i**, Oskar mediates Smaug localization and translational de-repression of *nanos* mRNA. With WT Oskar, Smaug, but not its co-factors in translational repression (Cup/CCR4-NOT), is localized to germ granules. Localized Smaug is dysfunctional in translational repression, allowing the translation of *nanos* mRNA. With Oskar-NQmut, Smaug loses the localization but remains functional inside germ granules, thus repressing the translation of *nanos* mRNA.

We reasoned that the selective localization of Smaug to germ granules should be controlled by particular granule protein components and might underlie the de-repression of *nanos* mRNA. Smaug has been shown to interact with Oskar, the scaffold protein that drives the assembly of germ granules and recruits mRNA ^35,38,39,71^. Oskar has an N-terminal LOTUS domain that mediates dimerization and binding to Vasa, a C-terminal SGNH-like domain with RNA binding function, and a 159-residue long linker region in between, which is predicted to be mainly intrinsically disordered (Fig. 4a) ^37,39^. Most of the *oskar* loss-of-function alleles identified so far have mutations within the LOTUS and SGNH-like domain and show defects in germ granule formation and RNA localization, precluding the analysis of the Oskar’s potential function as a translational regulator ^22,72^. The linker sequences of different *Drosophila* species are not conserved but enriched in the amino acids Asparagine (N) and Glutamine (Q) (Fig. 4b, and Extended Data Fig. 13a), which are over-represented in prion-like proteins ^73,74^. To probe the functional significance in these sequence features of Oskar in Smaug localization and *nanos* translation, we created a mutant Oskar protein where all the N and Q residues in the linker region were mutated to Glycine (*Oskar-NQmut*). We expressed the mutant protein at the anterior of the embryos via a *UA*S-*Oskar-NQmut-bcd3’UTR* transgene so that native germ granules at the posterior can serve as internal wildtype control (Fig. 4c). *Oskar-NQmut-bcd3’UTR* formed germ granules with indistinguishable morphology and localized *nanos* mRNA similar to Oskar-WT*-bcd3’UTR*. Furthermore, Oskar-NQmut granules exhibited similar physical properties as wildtype germ granules based on fluorescence recovery after photo-bleaching (FRAP) assay (Extended Data Fig. 13b, c). Thus, the N/Q residues in the Oskar linker region were not essential for mediating germ granule assembly, modulating material properties, or the recruitment of *nanos* mRNA. However, *Oskar-NQmut* germ granules completely lost their enrichment for Smaug (Fig. 4d, e), suggesting that the Oskar linker region mediates the recruitment of Smaug to germ granules, which is disrupted by N/Q-to-G mutations in the sequence.

Next, we investigated whether *Oskar-NQmut* affects *nanos* translation. To this end, we quantified the translation of *suntag-nanos* mRNA localized to the anterior *Oskar-WT* and *Oskar-NQmut* germ granules. We found a roughly 50% decrease in the percentage of translating *suntag-nanos* mRNA in germ granules composed of *Oskar-NQmut* protein compared to WT granules, suggesting a compromised translational function caused by this mutant (Fig. 4f, g). Consistent with reduced translation, the segmentation phenotypes caused by anteriorly expressed Nanos protein were much milder in *Oskar-NQmut-bcd3’UTR* embryos than in *OskarWT-bcd 3’UTR* embryos, validating that less Nanos protein was produced by *Oskar-NQmut* granules (Fig. 4h) ^29,35,47^. The reduction in translation could be due to a direct failure of Oskar protein to activate translation or to a loss of the ability to counteract Smaug-meditated repression in germ granules. We found that *suntag-nanos-SREmut* mRNA, which is not subject to repression by Smaug, was translated in *Oskar-NQmut* germ granules at a similar level as in WT germ granules (Fig. 4g), supporting the hypothesis that *Oskar-NQmut* granules are compromised in their ability to counteract the repression by Smaug. Together, these results suggest that Oskar controls *nanos* translation by mediating the selective sequestration of Smaug in germ granules (Fig. 4i).

## Discussion

It has been unclear whether biomolecular condensates can activate translation by directly serving as compartments for translation. Here, we utilized the single-molecule imaging method, SunTag, to visualize the translation of *nanos* mRNA *in vivo* to demonstrate that *Drosophila* germ granules are the sites for active translation while unlocalized mRNA is subject to translational repression. The SunTag system and high-resolution microscopy revealed the conformations adopted by translated and untranslated mRNA on germ granules. The quantitative nature of the SunTag system allowed us to dissect how germ granules affect translation efficiency and steady-state kinetics in vivo, which is not possible to unravel by conventional biochemical approaches. By mutating the disordered linker region of the scaffold protein Oskar, we uncovered its role in controlling the selective sequestration of translational repressors in germ granules, and, thereby, permitting *nanos* mRNA translation.

Our data distinguish *Drosophila* germ granules from most of the well-studied RNP granules that store translationally repressed mRNA and must be disassembled to resume mRNA translation. However, *Drosophila* germ granules might not represent a unique case of RNP granules providing space or a platform for translation ^9^. In fermenting yeast cells, mRNAs encoding glycolytic enzymes colocalize in specialized RNP granules and likely undergo translation within these granules ^19^. In the PGCs of zebrafish embryos, *nanos3* mRNA is suggested to be translated at the periphery of germ granules based on the distribution of ribosomes ^75^. *Pou5f3* mRNA granules in zebrafish embryos also have been shown to colocalize with nascent Pou5f3 peptides, suggesting the granules as translation sites ^18^. In mouse spermatids, liquid-liquid phase separation of FXR1 is essential for translational activation of FXR1 target mRNAs, suggesting that FXR1-RNA condensates are the compartments for activated translation ^17^. Interestingly, many of these examples were found in adult germ cells or early embryos, where transcription is largely inactive and translational regulation dictates the temporal and spatial distribution of proteins. Numerous specialized RNP granules or phase-separated condensates in germ cells and early embryos have been described so far ^76^. Thus, we expect more cases of translationally active RNP granules to be uncovered, establishing translational activation by RNP granules as a prevalent mechanism regulating gene expression.

High-resolution imaging allowed us to locate translating polysomes around the border or the surface of germ granules with 3’UTRs embedded internally. This extroverted orientation of polysomes is unlikely due to the lack of accessibility for translation machinery within RNP granules because we detected ribosomes or initiation factors inside granules. However, we propose that translation initiation, which requires sophisticated collaboration among multiple protein complexes, may be unfavored within a highly condensed environment ^77,78^. The correlation between translation status and the location of mRNA coding sequence suggests a potential regulatory mechanism: by controlling the inward or outward movement of a coding sequence, translation can be tuned up or down.

Notably, only 4% of total *nanos* mRNA localize to germ granules, while the remaining 96% is spread throughout the embryo’s soma ^31,52^. Inappropriate translation of *nanos* mRNA causes embryonic polarity and segment patterning defects ^29^. Therefore, strict translational repression of unlocalized mRNA and effective derepression by germ granules is necessary to establish the Nanos morphogen gradient emanating from the embryo posterior ^29,31,32,36,60,61,70^. Our imaging with *suntag-nanos* unambiguously demonstrates the repression-derepression dichotomy between soma and germplasm. Furthermore, we demonstrate that activation is achieved through increasing the fraction of translating mRNAs instead of alterations in ribosome occupancy or elongation rates. Our results are consistent with previous studies in tissue culture suggesting that the fraction of translating mRNA is highly variable among different mRNA and strongly affected by spatiotemporal regulation ^64^. Thus, controlling the translating fraction of an mRNA could be a common and critical aspect of translational regulation in vivo, and biomolecular condensates may be one mechanism to achieve this regulation.

Composition control plays an essential role in regulating the functions of biomolecular condensates. RNP granules often comprise a complicated set of RBPs, largely associated with translational repression ^79^. Consequently, RNP granules were considered translationally silent before rigorous tests using single-molecule imaging revealed some translation in stress granules ^69^. Our work shows that the enrichment of translational repressors does not necessarily render RNP granules translationally repressive but can instead underlie the translational de-repression mechanism of the target mRNA. Similarly, translation-activating condensates can form by RBPs that have long been considered as repressors such as FXR1 ^17,80^. Thus, the repression function of RBPs can be context dependent. Our results provide new insight into how repressors might become inactivated or altered when localized or assembled into condensates. It remains unclear how germ granule-localized Smaug loses its repressor function and how Oskar mediates this effect. We speculate that within germ granules, Smaug may lose interactions with cofactors like ME31B, Cup or CCR4 or change to a conformation that disfavors RNA binding, potentially via its specific interaction with Oskar’s linker region ^71^. Alternatively, interactions with germ granule proteins can constrain the mobility of Smaug and thus limit its access to the target mRNA *nanos* (Fig. 4i). Testing these hypotheses requires bottom-up reconstitution with purified proteins *in vitro* and characterization of germ granule-specific RBP interactomes *in vivo*.

## Acknowledgments

We thank M. Pamula, S. Grill, B. Lin, and J. Rajakumar for the constructive and critical feedback on the work and manuscript; E. Dawson for the initial conceptualization of the project. We thank P. Lasko and E. Wahle for sharing antibodies. We thank the Drosophila Genomics Resource Center for reagents, Bloomington Drosophila Stock Center and Kyoto Drosophila Stock Center for fly stocks.

## Author contributions

R.C. and R.L. designed the experiments. R.C., W.S., and J.D. performed the investigation. R.C. wrote the original draft. All authors reviewed and edited the text. R.L. and M.L. acquired the funding. R.L. supervised the work.

## Data and materials availability

All data needed to evaluate the conclusions in the paper are present in the paper and/or the Supplementary Materials.

## Supplementary information

Methods

Supplementary Notes

Extended Data Figs. 1 to 13

Tables S1 to S3

References

Movies S1 to S4

## Supplementary information

### Methods

#### Fly stocks

Fly stocks were maintained at 25°C. Detailed experimental genotypes and sources of fly stocks are listed in Table S1.

#### Cloning, gene-editing, and transgenesis

All primers are listed in Supplementary Table 2. All the constructs were made using In-Fusion cloning (Takara Bio) unless specified otherwise. All PCR was performed using CloneAmp HiFi PCR premix (Takara Bio #639298).

CRISPR: The SunTag array was knocked-in to the *nanos* locus by homology-directed recombination following CRISPR/Cas9 gene targeting ^1^. For generating the recombination template, the *nanos* sequence was PCR amplified from the genomic DNA of the *yw* Drosophila line. The SunTag sequence was PCR amplified from plasmid 5’TOP-SunTag-Renilla [Jeffrey Chao ^2^; Addgene #119946). The plasmid backbone and DsRed selection marker were PCR amplified from pScarless-HD-DsRed (Kate O’Connor-Giles, Addgene #64703) and assembled with *nanos* and SunTag fragments using In-Fusion assembly. SunTag array was placed right after the start codon of *nanos* open reading frame while the DsRed marker was inserted after a TTAA sequence of the first intron of *nanos* to allow transposase-mediated excision. Two guide RNAs (guide #1: GATAACCGTAACTTTCGACC; guide #2: GTAAGAAGAAATGGCGAATA) were separately cloned into pCFD-dU6:3gRNA (DGRC_1362) by linearizing the plasmid with primers appended with the guide RNA sequences and ligation with KLD enzyme mix (NEB M0554S). The recombination template and two guide RNA plasmids were injected into various Cas9-expressing lines (BestGene Inc.). Transformant flies were screened using the DsRed eye marker. Also, see supplementary text for details about the generation of the *suntag-nanos* CRISPR line.

##### UAS-suntag-nanos transgenes

To generate the *UAS-suntag-nanos* construct, the *nanos* sequence starting from the 5’ UTR to 500 nucleotides following the end of the 3’UTR was PCR amplified from genomic DNA of *yw* strain and inserted into PCR-linearized pUASz1.1 plasmid [Allan Spradling ^3^; DGRC_1433), via In-Fusion assembly. The 5’UTR of *nanos* is placed right after the Hs promoter of the plasmid so that IVS and Syn21 elements of the pUASz1.1 plasmid are removed. The P10 3’UTR of the pUASz1.1 plasmid was also left out during cloning. The SunTag sequence was amplified from 5’TOP-SunTag-Renilla (Addgene #119946) and inserted after the *nanos* start codon via In-Fusion assembly.

To introduce mutations at the two SRE sequences in the *nanos* 3’UTR ^4,5^, a gBlock of the sequence containing the mutant SREs was synthesized and replaced the WT sequence in the *UAS-suntag-nanos* plasmid. The wild-type sequences are SRE1: GCAGAGGCT<u>C</u>T<u>G</u>GCAGCTTTTGC, and SRE2: AAATAGCGC<u>C</u>T<u>G</u>GCGCGTTCGAT. The mutant has underlined C mutated to G, and underlined G mutated to C)

The sequence of tubulin84B 3’UTR was amplified from genomic DNA and replaced the *nanos* 3’UTR of *UAS-suntag-nanos* plasmid to generate *UAS-suntag-nanos-tub3’UTR* ^6^.

##### UAS-osk-bcd3’UTR transgenes

Full-length oskar coding sequence and 3’UTR of *bicoid (bcd 3’UTR)* without the Nanos-response element (NRE) were PCR amplified from plasmid UAS-oskar-mCherry3xFLAGHA-bcd3’UTR ^7^. pUASz1.1 vector was PCR-linearized and assembled with *oskar* CDS and *bcd 3’UTR* fragment to generate *UAS-osk-bcd 3’UTR*. To generate *oskar-NQmut*, a gBlock of Oskar linker region containing N/Q-to-G mutations was synthesized and replaced the WT sequence of *UAS-osk-bcd 3’UTR* plasmid.

For transgenesis, individual plasmids were injected into attP2 or attP40 lines (the BestGene Inc.) and transformants were screened for presence of the DsRed eye markers.

#### Immunofluorescence (IF)

Drosophila embryos were collected for 3h on an apple juice plate, dechorionated by incubating with 50% bleach solution for two minutes, extensively washed, and transferred to a scintillation vial containing a 1:1 (v/v) mixture of heptane and 4% paraformaldehyde in PBS (phosphate-buffered saline), in which embryos were permeabilized and fixed for 20 min. The paraformaldehyde was removed with a Pasteur pipette, followed by adding methanol and vigorous shaking for 15 s to remove the vitelline membrane. Embryos were washed three times for 5 min with methanol before being stored at 4°C in methanol. Embryos were rehydrated by washing for 5 min with 50% methanol with PBS-Triton 0.3% and then washed and permeabilized for 3x 15 min in PBS-TritonX-100 0.3%. Embryos were blocked with 1% BSA in PBS-TritonX-100 0.3% for 30 min and subsequently incubated with primary antibodies diluted in the blocking solution (anti-Oskar (gift from Paul Lasko) 1:1000, rabbit anti-Nanos (Lehmann Lab) 1:1000, rabbit anti-CCR4 and anti-NOT3 (gift from Elmar Wahle) 1:1000, rabbit anti-RPS6 (Cell Signaling #2217) 1:200, rabbit anti-GCN4 (Novus Bio, clone C11L34) 1:1000) overnight at 4°C. Embryos were washed five times for 10min with PBS-TritonX-100 0.3%, blocked for 30min, and incubated with secondary antibodies (ThermoFisher Scientific anti-rabbit AlexaFluor488, anti-rabbit AlexaFluor647, anti-rat AlexaFluor555) with 1:1000 dilution for 4 h at room temperature. Then embryos were washed five times for 10 min with PBS-Triton 0.3%, stained with DAPI, and mounted with ProLong Glass mounting medium (ThermoFisher, P36980).

#### Single-molecule fluorescence in situ hybridization (smFISH)

The smFISH with fixed embryos and ovaries is modified based on ^8,9^. Stellaris RNA FISH probes against *suntag*, *nanos*, and *oskar* sequences were used for hybridization. The *nanos* CalFluor 590 and *oskar* CalFluor 590 probes have been described and used in the previous work ^10^. The *suntag* Quasar 670 and *nanos 3’UTR* Quasar 670 probes were synthesized by LGC Biosearch Technologies. The probe sequences are listed in Supplementary Table 3. To perform smFISH on fixed embryos, stored embryos were rehydrated by washing for 5 minutes with 50% methanol with PBS-Tween 0.1% and washing three times for 5 min in PBS-Tween 0.1%. Embryos were then washed with pre-hybridization buffer containing 2xSSC and 10% formamide (Fisher Scientific, AM9342) for 10 minutes at room temperature. The embryos were then incubated at 37°C for 3h in the hybridization mix (60µL hybridization mix per sample with 50-100 embryos) containing 2xSSC, 10% (v/v) deionized formamide, 0.1 mg/ml E.coli tRNA, 0.1mg/ml salmon sperm DNA, 10mM Ribonucleoside Vanadyl Complex (NEB, S1402S), 2mg/ml BSA, 80ng Stellaris probes and 10% (v/v) Dextran sulfate. After hybridization, embryos were washed with pre-hybridization buffer twice for 15 minutes at 37°C. The embryos were washed with PBS-Tween 0.1% three times for 5min, stained with DAPI, and mounted with ProLong Glass mounting medium.

When anti-GCN4 is used to detect SunTag protein, IF was performed after smFISH. Following the 2×15min washes with pre-hybridization buffer, embryos were washed and permeabilized with PBS-TritonX-100 0.3% for 45 minutes. Embryos were blocked with 1% BSA in PBS-TritonX-100 0.3% for 30 minutes and then incubated with rabbit anti-GCN4 (Novus Bio.) with 1:1000 dilution overnight at 4°C. Embryos were washed with PBS-TritonX-100 0.3% five times for 10 minutes, blocked for 30min, and incubated with anti-rabbit secondary antibody (1:1000) for 4h at room temperature. Embryos were then washed with PBS-Triton 0.3% five times for 10 min, stained with DAPI, and mounted with ProLong Glass mounting medium.

To detect the translation of *suntag-nanos* in ovaries, *Vasa-mApple/+; suntag-nanos, scFv-GFP/+* flies were used and ovaries were hybridized with *suntag* smFISH probes. Ovaries were dissected out in Robb’s buffer (100 mM HEPES, 100 mM sucrose, 55 mM sodium acetate, 40 mM potassium acetate, 10 mM glucose, 1.2 mM magnesium chloride, and 1 mM calcium chloride) and fixed with 4% paraformaldehyde in Robb’s buffer for 20min. After fixation, ovaries were washed with PBS-Triton 0.3% twice for 5min, 50% methanol in PBS-TritonX-100 0.3% for 5min, 100% methanol for 30min. Ovaries can be stored in methanol at 4°C. The rehydration, hybridization, and washing for smFISH follow the same protocol as for embryo samples described above.

The detailed genotypes of flies and reagents for fluorescence labeling (IF/smFISH) used in each experiment are listed in Supplementary Table 1. Specifically, to detect *suntag-nanos* mRNA using probes against *nanos* 3’UTR, embryos from Vasa-mApple/+; *suntag-nanos, scFv-GFP/Df(3R)DlSP* flies are used, where *Df(3R)DlSP* is a deficiency line that does not contain the *nanos* locus and thus *suntag-nanos* allele is the only source of the *nanos* mRNA in embryos (*30*). For experiments involving *UAS-suntag-nanos-SREmut, UASz-suntag-nanos-tub3’UTR,* and *UAS-oskWT/NQmut-bcd3’UTR,* which increased *suntag-nanos* translation, we used anti-GCN4 to detect SunTag instead of scFv-GFP due to a potential depletion of scFv-GFP in the embryos (*45*).

#### Confocal microscopy of fixed embryo samples

Images of whole embryos were acquired using Zeiss LSM780 confocal microscope with 10x 0.3 Numerical Aperture (NA) air objective and 2.2 pixels per micron. Red fluorophores (mApple, mCherry, Alexa Fluor 555, or Alexa Fluor 561) were excited by a 561nm laser. Green fluorophores (GFP, YFP, or Alexa Fluor 488) were excited by a 488nm laser. The far-red fluorophore (Alexa Fluor 647) was excited by a 633nm laser.

High-resolution images of germplasm were acquired using Zeiss LSM980 confocal microscope with Plan-Apochromat 63x /1.4NA oil objective with AiryScan 2 detector and SR mode. GFP and Alexa Fluor 488 were excited using a 488nm laser; mApple, mCherry, Alexa Fluor 555, and Alexa Fluor 568 were excited using a 561nm laser; Alexa Fluor 647 and Quasar 670 were excited using a 639nm laser. Images were acquired with 1.7x zoom, 23.5 pixels per micron, 8 bits per pixel, and without averaging (result imaging size 78.2 µm x 78.2 µm, 1840 pixels x 1840 pixels). Multiple z-stacks (10∼30 stacks) were taken with a 150 nm interval (Voxel size: 42.5 nm x 42.5 nm x 150 nm). Raw images were first processed with the 3D AiryScan processing function in ZEN Blue software. Imaging TetraSpeck™ Fluorescent Microspheres showed clear chromatic aberrations among three channels. Therefore, aberration correction files were generated using the channel alignment function in ZEN by correcting the signal misalignments in a microsphere image. The correction files were applied to correct the embryo images (post-Airyscan processing) using the channel alignment function. The images after Airyscan processing and channel alignment were saved as final data and used for later analysis and publication.

#### Image analysis of translation foci and quantification

SunTag image analysis was performed using MATLAB-based software FISH-Quant_v3, which allows the detection of focal signals and analysis co-localization in a three-dimensional space (3D) ^11^. Images taken from Zeiss LSM980 using the 63x oil lens and AiryScan 2 detector with three channels (germ granules marked by Vasa, *suntag* mRNA smFISH, SunTag protein stained by scFv-GFP or anti-GCN4) were first split using Fiji software. The Vasa channel images were used to define the outline of germplasm and soma. To detect the foci of *suntag* mRNA and SunTag protein (anti-GCN4/scFv-GFP), a pre-detection was performed to test a range of threshold values and determine the number of detected spots for each tested value. The number of detected spots usually plateau at a range of tested thresholds and the number increased exponentially with lower thresholds which indicated the detection of background or noise signals. The threshold was placed at the left side of the plateau range before the increase occurred and foci were detected with this set threshold. The detected spots were then fit with a 3D Gaussian, which determined the 3D coordinates and intensities of individual foci for later analysis. The co-localization between the detected mRNA and protein foci was analyzed by the DualColor program of FISH-Quant. We set 400 nm as the maximum distance between two spots to be considered co-localized although the number of co-localization events usually plateaus at 250nm. This analysis provides the percentage of mRNA foci co-localized with protein foci, which represents the percentage of translating mRNA.

#### Distance measurement

To measure distances between the mRNA and SunTag foci and the germ granule surface, images taken from Zeiss LSM980 using 63x oil lens and AiryScan 2 detector with three channels (germ granules marked by Vasa-mApple or Vasa-GFP, *suntag-nanos* mRNA stained by smFISH probes against *suntag* or *nanos 3’UTR*, SunTag protein stained by scFv-GFP or anti-GCN4) were used. Images were first analyzed with FISH-Quant_v3 and DualColor as described in the section above, which provides a result file with the *x,y,z* coordinates of mRNA and SunTag foci and identifies translating and non-translating mRNA.

The machine learning-based image analysis program *ilastik* was used to segment germ granules visualized in the Vasa channel ^12^. The classifier in the Pixel Classification workflow was trained using 3 representative images and was then applied to unseen images to perform binary segmentation of germ granules. The segmented germ granule files were then imported into FIJI and analyzed using a FIJI macro. Briefly, this macro outputs files with the coordinates of each pixel categorized as belonging to germ granules based on the previous ilastik-based segmentation. Various pixel lists were compiled, which separated the pixels on the 3D surface of the granule (identified using the 3D Object Counter plugin in FIJI) from those entirely within the granule. Additionally, only the granules that were entirely within the acquired image (in all three dimensions) were used in the analysis. In other words, the granules that contacted the x,y, or z border of the image were identified and excluded from the analysis because the borders of the image provide artificial surfaces for the granules and may affect the outcome of the analysis.

The 3D segmentation of germ granules and coordinates of SunTag and mRNA foci acquired from FISH-Quant analysis were then further analyzed in a custom-built Python workflow to perform the 3D distance measurement of mRNA and SunTag foci from the surface of the germ granule. First, the minimum distance of each point to the closest pixel on the 3D surface of the granule was calculated. Next, any mRNA or scFv foci that were closest to a granule that touched the edge of the Z-stack were excluded. This step was performed to ensure that any analysis on localization was only performed on foci associated with granules that were fully captured within the image. Then, the foci were categorized as being inside or outside the granule based on their relative position to the 3D surface pixels of the granule compared to pixels entirely within (or outside) the granule. After the categorization of foci as either inside or outside granules, the minimum distance to the surface of the germ granule values were adjusted to be negative if the foci were inside the granule and kept as positive if the foci were outside the germ granule. The adjusted distance values were then plotted as relative frequency histogram using Seaborn in Python. A bin size of 25 nm was used. Kernel density estimate (KDE) plots were generated and overlaid with the histogram.

#### Computation controls of 3D distance measurements

A representative section of germplasm with relatively even coverage of germ granules across the image was first cropped from an image. The 3D distances of mRNA foci from the surface of the germ granules in this image were obtained and plotted as detailed in “3D Distance Measurements of mRNA and SunTag foci from germ granule surface”. Then, the mRNA channel of the image was rotated 180° and analyzed and plotted using the same workflow. Next, 1 million points (located within the volume of the image) were generated by drawing from a uniform distribution in x, y, and z. These simulated points were then analyzed using the same workflow previously mentioned.

#### Puromycin injection

Embryos from *Vasa-mApple/+; suntag-nanos, scFv-GFP/+* flies were dechorionated using 50% bleach for 2 min and washed thoroughly with water. About 40∼50 embryos were then lined up and mounted at the edge of a coverslip by heptane glue with their posterior poles pointing toward the edge of the coverslip. Mounted embryos were placed in a desiccator at 18°C for 10 min and then covered by halocarbon 700 oil. 20mM HEPES (control) or 10mg/ml puromycin in 20mM HEPES (Gibco, A1113803) was injected at the posterior pole of the embryos using FemtoJet (Eppendorf, 5252000021) with Femtotips II (Eppendorf, 930000043) needles at 18°C. The exact injected volume of solution was difficult to control but generally, the volume was small to avoid pushing cytoplasm out of embryos. The injected embryos were aged for 15 min to 30 min at 18°C before being transferred to a glass vial containing a 1:1 (v:v) mixture of heptane and fixative (4% paraformaldehyde in PBS) and fixed for 60min at room temperature. After fixation, embryos were transferred onto a double-sided tape within a petri dish and covered with PBS-Tween 0.1%. The vitelline membrane was removed manually with the needle of an insulin syringe. Devitellinated embryos were stepped into 100% methanol and washed in methanol three times for 5min before being stored or proceeding to smFISH.

#### Live imaging and FRAP

We found that fertilized embryos showed apparent cytoplasmic movement during live imaging, potentially due to the mechanical force generated during nuclear division and migration ^13^. We found that unfertilized eggs had significantly less cytoplasmic movement, which allowed tracking individual polysomes for extended periods (>5 min) and thus were used for the live imaging.

For live imaging of *suntag-nanos* mRNA translation, we expressed *UAS-suntag-nos* (WT or SREmut) with a weak Matα-GAL4 (without VP16) maternal driver line to prevent scFv-GFP depletion which can cause artifact (Extended Data Fig. 2) ^14,15^. Unfertilized embryos were collected from *Vasa-mApple/Matα-GAL4; uas-suntag-nos (WT or SREmut)/scFv-GFP* virgin female flies that mated with sterile male flies (male progeny of osk301/oskCE4 flies). Dechorionated embryos were mounted onto the coverslip of a glass bottom 35mm dish with their posterior poles pointed toward and glued onto the coverslip (Fig 3d) to allow the best imaging of germplasm. Live embryos were imaged on Zeiss LSM980 confocal microscope through the 63x oil objective lens (Plan-Apochromat, 1.4 NA) using AiryScan 2 detector and SR mode. Germplasm was first located and moved into focus using Vasa-mApple through a red fluorescence channel (excitation laser 561nm) with 1x zoom. Then a small region-of-interest (ROI) was imaged (292 pixels x 292 pixels, 12.45 µm x 12.45 µm) and translation sites (bright GFP foci) were identified through GFP channel (excitation laser 488nm). Time-lapse images (movies) were acquired with 10 seconds per frame and 40 frames in total. In each frame, 25 z-stacks with a 150nm interval were imaged with the GFP channel only. For the FRAP experiment, multiple regions containing translation sites were selected and photo-bleached with 70% power 488nm laser for 10 iterations. Three frames were taken before the bleaching.

Images were analyzed as maximum intensity projections. We tracked individual translation sites with a Fiji plugin, TrackMate (v6.0.2). LoG detector was used to detect translation sites, with an estimated blob diameter 0.4µm and sub-pixel localization. Detection thresholds were adjusted for individual images. Simple LAP tracker was used to track the foci movement, with a maximum gap distance of 1 µm and a maximum gap of 1 frame. Although overall cytoplasmic movement is reduced in unfertilized eggs, translation sites were still undergoing constant and stochastic movement and might move out of the imaging field which resulted in most of the short tracks in the tracking result. Therefore, only long tracks (>30 frames, 5 minutes in total) were selected for further analysis. For the photo-bleached GFP foci, they were usually undetectable by the tracking program for 4-6 frames before fluorescence recovered to the detection limit. During this period, photo-bleached foci were manually tracked; the tracks of the same spot before bleaching and after fluorescence recovery could be manually identified and connected. The intensities of individual foci in each frame were then extracted from the result file and plotted with time on Prism 8.

#### Analysis of FRAP data and calculation of translation elongation rate

The basic assumptions of using FRAP experiments to calculate translation elongation rate have been discussed previously ^16–18^. We assume that 1) ribosomes are uniformly distributed within the open reading frame (ORF); 2) ribosome elongation rate is constant; 3) scFv-GCN4 epitope binding is stable and exchange at a significantly slower rate than translational elongation (*44*); 4) nascent peptides are immediately released when synthesis completes.

The first phase of fluorescence recovery after the photo-bleaching is linear due to the synthesis of new SunTags at a constant rate while the fully-synthesized and released SunTag-Nanos proteins were still labeled with photo-bleached scFv-GFPs, thus not contributing to the signal change of the polysome. We defined that L1 is the length of the SunTag array and L2 is the length of the Nanos. The linear phase lasts until the fluorescence intensity recovers to *I*_0_ × *L*_2_/(0.5*L*_1_ + *L*_2_), where *I0* is the initial fluorescence intensity before bleaching.

And the recovered fluorescence intensity over time: 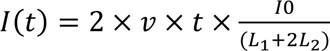

In the equation, *t* is the time after photo-bleaching and *v* is the elongation rate.

The second phase starts when the first SunTag synthesized post-photo-bleaching leaves the polysome with fully-synthesized protein, which counteracts the increase of newly synthesized SunTag and causes the increase of signal to slow down. When the first ribosome loaded after photo-bleaching finish the translation, the signal reaches a plateau (the third phase) with the same intensity as before photo-bleaching because all the SunTags are bound by non-bleached scFv-GFP again. Indeed, our FRAP curves showed these three phases (Extended Data Fig. 9c). We used the data of the linear phase to fit the linear equation above to calculate the translational elongation rate.

#### Calculation of ribosome occupancy

The rationale and mathematical basis of calculating ribosome occupancy using fluorescence of polysomes and single SunTag protein is based on previous studies ^16–21^. To measure ribosome occupancy, we used rabbit anti-GCN4 antibody to detect the SunTag peptides in embryos from *Vasa-mApple; suntag-nanos* flies, which provides high signal-to-noise ratio and allows clear visualization of single fully-synthesized SunTag-Nanos protein (Extended Data Fig. 10a). Fluorescence intensities of polysomes are generally five to ten-fold higher than a single synthesized SunTag-Nanos peptide so polysomes can be detected in FISH-Quant without detecting single peptides by setting a relatively high threshold. In fact, the automatically assigned threshold in the pre-detection step in FISH-Quant has always been higher than single peptides. To specifically detect single peptides with FISH-Quant, a region in soma where there are only single peptides (no polysome) is selected, and a low threshold is used for pre-detection (at least ten-fold lower than the automatically assigned threshold). In the pre-detection plot, a slope is usually observed at a range of low thresholds before the exponential increase of the detected number at lower thresholds, which corresponds to the background signal. The detection threshold is placed in the middle of the slope, which can capture most of the distinguishable single peptide spots while leaving out the dimer spots which may represent the degrading peptides or peptides not fully labeled. This may also cause an overestimation of the intensity of a single SunTag protein and consequently under-estimation of ribosome occupancy. Raw intensities of polysomes and single peptides were extracted from the result file of FISH-Quant analyzed, and plotted in Prism to obtain mean intensities of each population: *F* = intensity of a polysome, and *F0* = intensity of a single protein.

As the ribosomes within the SunTag CDS only synthesize part of the SunTag repeats, they don’t contribute to the fluorescence as much as the ribosomes within the Nanos CDS, which have the complete SunTag repeats. Assuming ribosomes are uniformly distributed throughout the CDS, the ribosomes within the SunTag repeats on average have half of the SunTag repeats. Therefore, the effective length of the open reading frame equals 0.5*L*_1_ + *L*_2_, in which *L2* is the length of Nanos and *L1* is the length of the SunTag. The fluorescence intensity of a polysome *F* = *F*_0_ × *d* × (*L*_2_ + 0.5*L*_1_), in which *d* is the density of ribosomes in a polysome.

#### *In vitro* egg activation

The protocol is adapted based on previous studies ^22,23^. Young (less than one week old) *Vasa-mApple/+; suntag-nanos, scFv-GFP/+* flies were well fed to enrich late-stage oocytes, which were then dissected out in 1x Robb’s buffer. Stage 14 oocytes were identified and sorted out based on the morphology of the dorsal appendages and transferred into 30% Robb’s buffer to be incubated and activated for over 30min, during which oocytes became swollen and some dorsal appendages became separated. Incubated oocytes were incubated in 50% bleach for 1 min, during which nonactivated oocytes were lysed by bleach while activated oocytes survived due to vitelline membrane cross-linking and were immediately washed thoroughly with 30% Robb’s buffer. Activated oocytes were mounted on the coverslip of a glass bottom dish with the posterior pole stuck onto the coverslip by heptane glue. A small piece of wet tissue was put inside the dish to humidify the internal. Oocytes were imaged live with Zeiss LSM980 confocal microscope with the 63x oil lens and Airyscan 2 detector.

#### Oskar sequence feature analysis

Sequences of 11 Drosophila species (*D.melanogaster, D.immigrans, D.virilis, D.hydei, D.miranda, D.grimshawi, D.navojoa, D.pseudoobscura, D.arizonae, D.persimilis*) were aligned and conservation plot was acquired in Benchling. Disorder sequence prediction was performed in IUPred2A website (https://iupred2a.elte.hu/) ^24^.

#### Quantification of translation in germplasm induced by *Oskar-NQmut*

To compare the *suntag-nanos* translation on Oskar-NQmut germ granules with WT Oskar, we generated flies expressing *suntag-nanos* or *suntag-nanos-SREmut*, together with *UAS-Oskar-WT/NQmut-bcd3’UTR* transgene to induce germ granules at the anterior pole of the embryos (detailed genotypes in supplementary table 1). Embryos were collected from the flies, fixed and stained with suntag smFISH probes and anti-GCN4. Images were acquired from the induced germplasm at the anterior pole as well as the native germplasm at the posterior. The percentage of translating mRNA was measured using FISH-Quant using the quantification protocol described above. The translation activity at the anterior of an embryo is measured by the translating fraction of the anterior germplasm normalized with the fraction of the posterior germplasm.

#### FRAP of germ granules

To assess the dynamics of germ granules made by Oskar-WT or Oskar-NQmut, embryos expressing Vasa-mApple and *Oskar-WT-bcd3’UTR* or *Oskar-NQmut-bcd3’UTR* were collected, mounted with anterior poles on coverslip, and imaged live with Zeiss LSM780 with red fluorescence channel (excitation wavelength 561 nm). An area of about 5 µm × 5 µm size was chosen in germplasm and photo-bleached with 70% 561nm laser. The fluorescence of the bleached region was recorded with time-lapse imaging (5s interval), measured as integrated intensity using Fiji, and plotted over time using Prism 8.

#### Cuticle prep and imaging

Embryos were collected overnight from the *Matα-GAL4VP16/UAS-osk (WT or NQmut)-bcd3’UTR* flies and aged for 24 hours, after which embryos were dechorionated with 50% bleach for 2 min, extensively washed, and then transferred to a mesh-bottom basket. The embryos in the basket were incubated with the acidic acid/glycerol 4:1 mixture for 1 hour at 60°C, after which embryos were transferred to a slide, covered with Hoyer’s medium and coverslip, and incubated overnight at 60°C. The cuticles were examined with a dark field stereomicroscope.

#### Supplementary Notes

##### Generation of CRISPR *suntag-nanos* line

We made a substantial attempt to knock-in SunTag array into the *nanos* locus using CRISPR-Cas9 genome editing. With the repair template and the two guide RNAs described in the method section, over 2000 embryos were injected by *BestGene*. Only one transformant was identified in the screen, which was used to establish the *suntag-nanos* line. In the repair template, the SunTag array was placed after the start codon of the Nanos open reading frame (ORF), and a DsRed marker cassette flanked by P-Bac transposon ends was put into the first intron of the *nanos* gene, which could be subsequently removed by P-Bac transposase. The sequencing of genomic DNA of the *suntag-nanos* line showed that the SunTag array was inserted into the correct position in the Nanos ORF. The DsRed marker cassette, however, was not in the first intron of *nanos*. Instead, the cassette, together with some flanking *nanos* sequences (parts of the left and right homology arms), was inserted upstream of *nanos* locus (about 300bp upstream of the transcription start site). This unexpected insertion may be the result of a rare splicing event of the repair template during homology-directed repair. In addition, we used different guide RNAs and/or different repair template designs (not presented in this paper), attempting to generate more transformant, but without success. As the SunTag was correctly inserted into the designed position in the *suntag-nanos* line, and our experiment validated that the *suntag-nanos* mRNA showed similar RNA localization and translation pattern as native *nanos* mRNA, we used this line throughout our study.

##### Quantification of *suntag-nanos* mRNA translation

The *suntag-nanos* line is homozygous-viable. Female flies homozygous for the *suntag-nanos* insertion layed a similar number of eggs as wildtype flies but the embryos did not hatch. This suggests that SunTag-Nanos can perform the function of Nanos in germline stem cells but is unable to support posterior patterning in embryos. The insertion of the DsRed marker upstream of the *nanos* locus might disrupt an enhancer for *nanos* transcription because the mRNA abundance of the *suntag-nanos* RNA was significantly lower than that of the native *nanos* gene. The low expression, however, was advantageous for the quantification of the translating fraction. Under normal expression levels, *nanos* mRNAs form homotypic clusters, whereby each cluster contains multiple copies of *nanos* mRNA per germ granule. Thus, distinguishing between translating and non-translating mRNA using SunTag in a multi-copy mRNA cluster would have been technically challenging. The low expression of *suntag-nanos* reduces *nanos* mRNA levels to, on average, one mRNA per granule. This allowed us to quantify the translation of mostly single *suntag-nanos* mRNA per granule.

**Extended Data Fig. 1.**
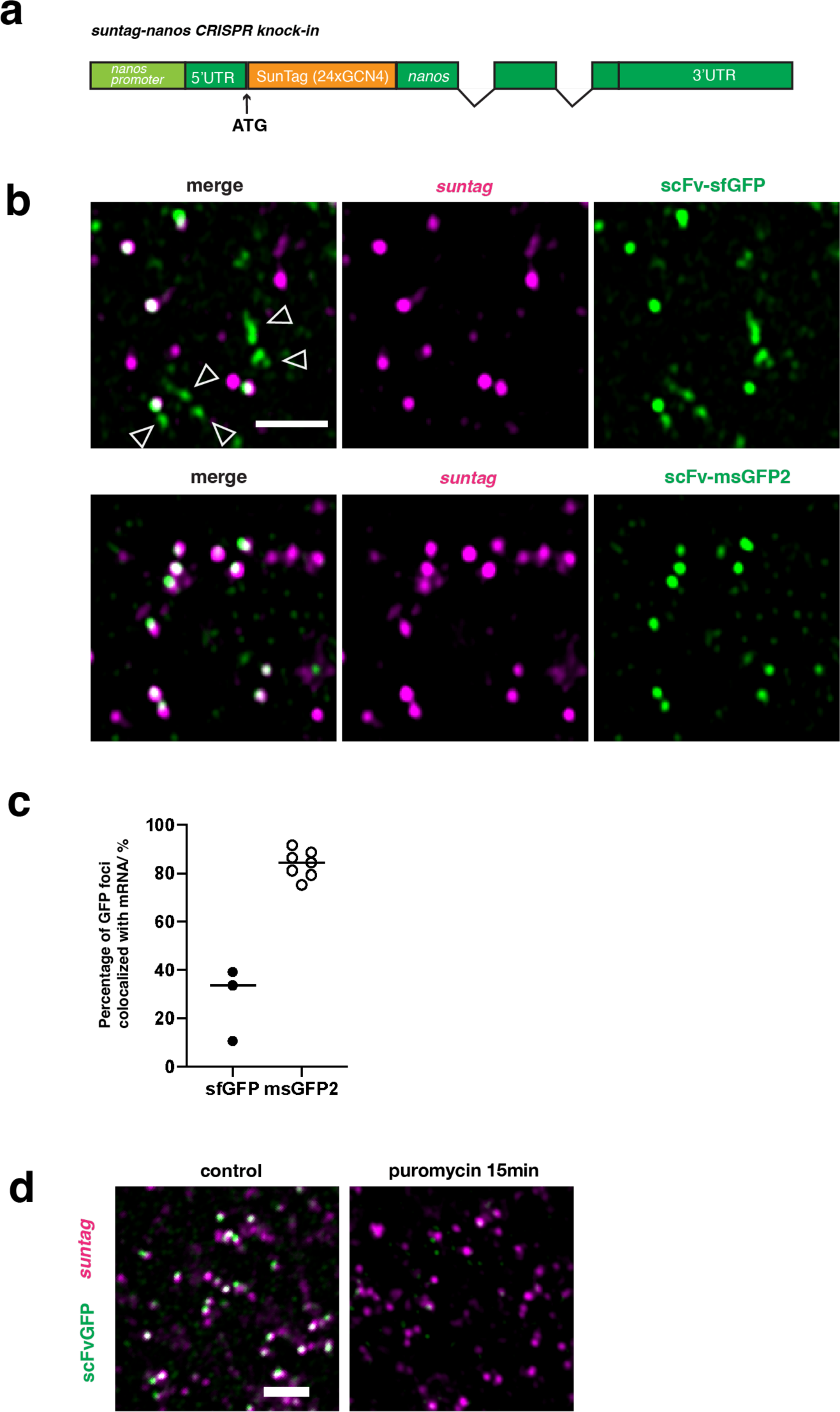
Optimization and validation of the *suntag-nanos* system. **a**, Schematic of CRISPR knocked-in *suntag-nanos* allele. **b**, Images of germplasm in embryos expressing *suntag-nanos* and (top) scFv-sfGFP (super-folder GFP) or (bottom) monomeric msGFP2 (green). *Suntag* mRNA is stained by *suntag* probes (magenta). ScFv-sfGFP showed puncta GFP signals (arrowheads) which are not co-localized with mRNA signal and thus are not translating sites. ScFv-msGFP2, which is used throughout this study unless suggested otherwise, significantly reduces the aggregation. Scale bar 2 µm. **c**, The percentage of GFP foci co-localized with mRNA. The aggregate formation of scFv-sfGFP causes a relatively low percentage of colocalization. Using scFv-msGFP2, the majority of GFP foci (80%-90%) represent polysomes. **d**, Images of germplasm in embryos expressing *suntag-nanos* and scFv-GFP (green). Embryos were injected with 20 mM HEPES or 10 mg/ml puromycin and aged for 15min before being stained with *suntag* probes (magenta). Scale bar 2 µm.

**Extended Data Fig. 2.**
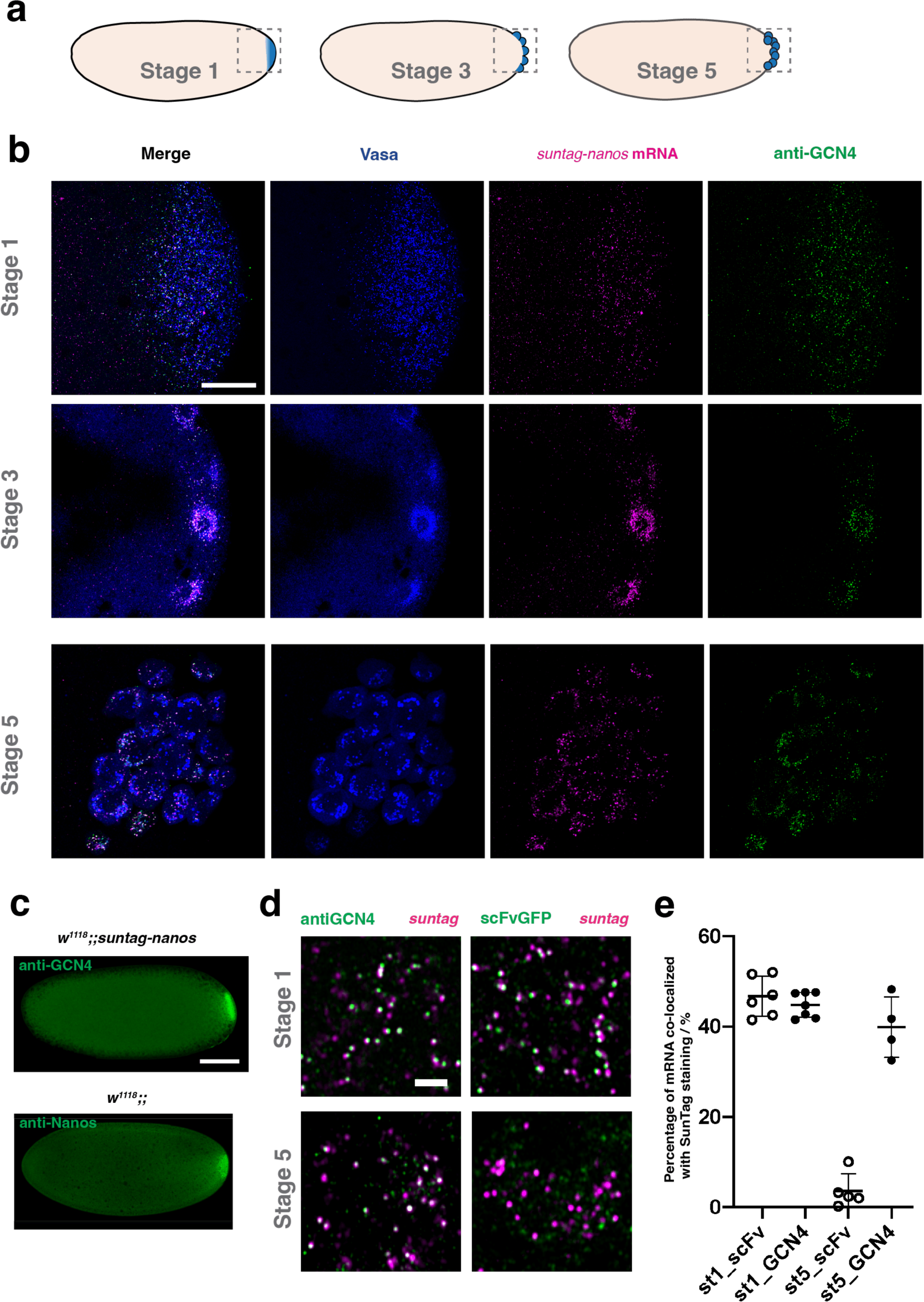
Detecting SunTag with anti-GCN4. **a**, Schematics of stage-1, stage-3, and stage-5 embryos. Germplasm or pole cells are labeled in blue. Outlined regions are imaged and presented in panel (B). **b**, Example images of stage-1, stage-3, and stage-5 embryos expressing *suntag-nanos*. SunTag is stained by anti-GCN4 (green); suntag mRNA is stained by smFISH (magenta); germplasm or pole cells are marked by Vasa-mApple (blue). Scale bar 20 µm. **c**, Images of a stage-1 embryo expressing *suntag-nanos* stained by anti-GCN4 (top) and a stage-1 embryo from a *w*^1118^ fly stained by anti-Nanos (bottom). Scale bar 100 µm. **d**, Images of germplasm in stage-1 embryos (top) and pole cells in stage-5 embryos expressing *suntag-nanos*. SunTag (green) is stained by anti-GCN4 (left) or by endogenous scFv-GFP (right); *suntag* mRNA is stained by smFISH (magenta). Scale bar 2 µm. **e**, Quantification of the percentage of mRNA foci co-localized with SunTag staining signal in stage-1 germplasm and stage-5 pole cells when SunTag is stained by scFv-GFP or anti-GCN4.

**Extended Data Fig. 3.**
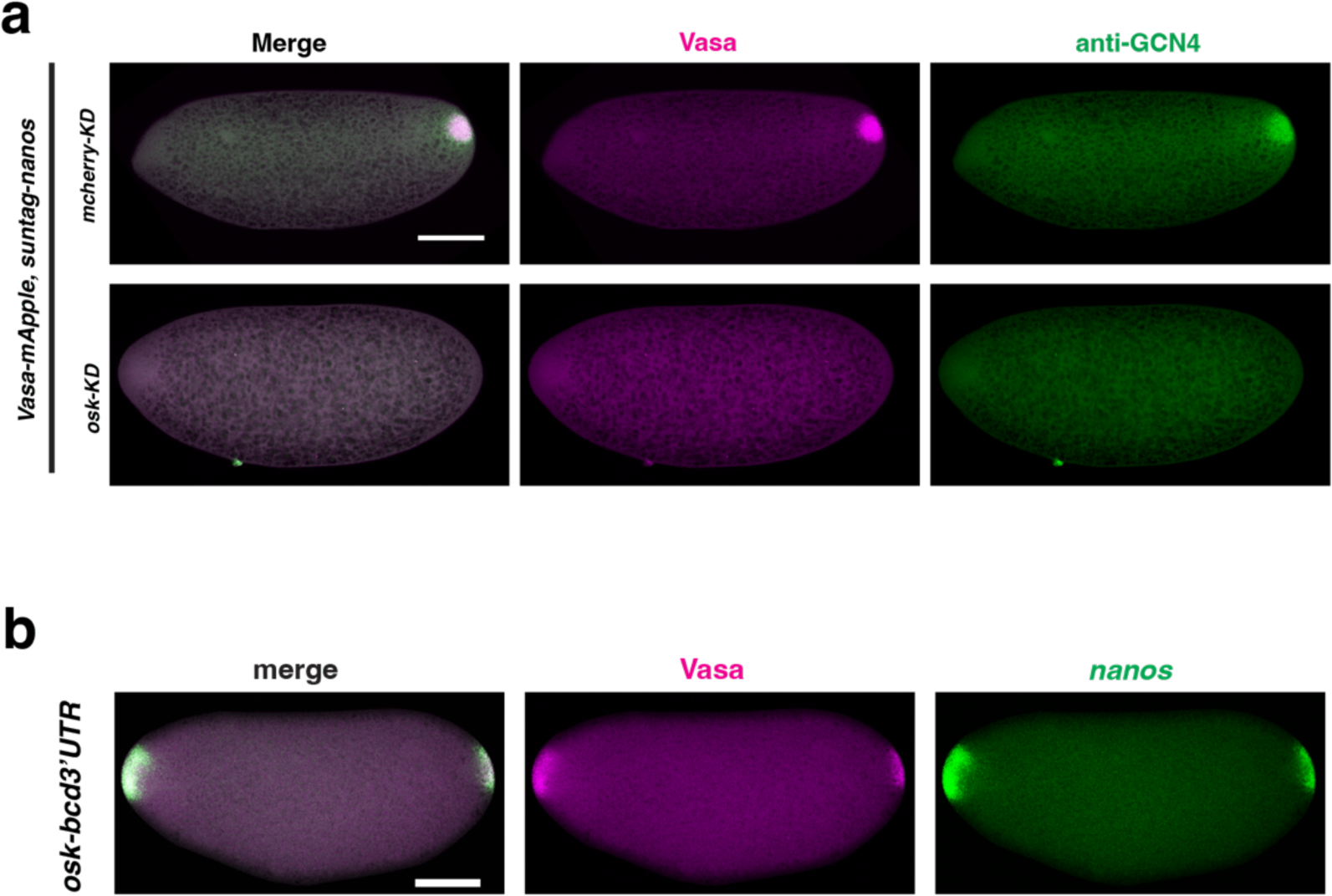
*Suntag-nanos* mRNA translation depends on germplasm. **a**, Images of embryos expressing *suntag-nanos* with *mcherry* knockdown (top) or *osk* knockdown (bottom). SunTag is stained by anti-GCN4 (green) and germplasm is marked by Vasa-mApple (magenta). Scale bar 100 µm. **b**, Images of embryos expressing Vasa-mApple and *osk-bcd3’UTR*, forming germplasm and localizing *nanos* mRNA at the anterior pole. Germplasm is marked by Vasa-mApple (magenta). Endogenous *nanos* mRNA is stained by smFISH probes against *nanos* (green). Scale bar 100 µm.

**Extended Data Fig. 4.**
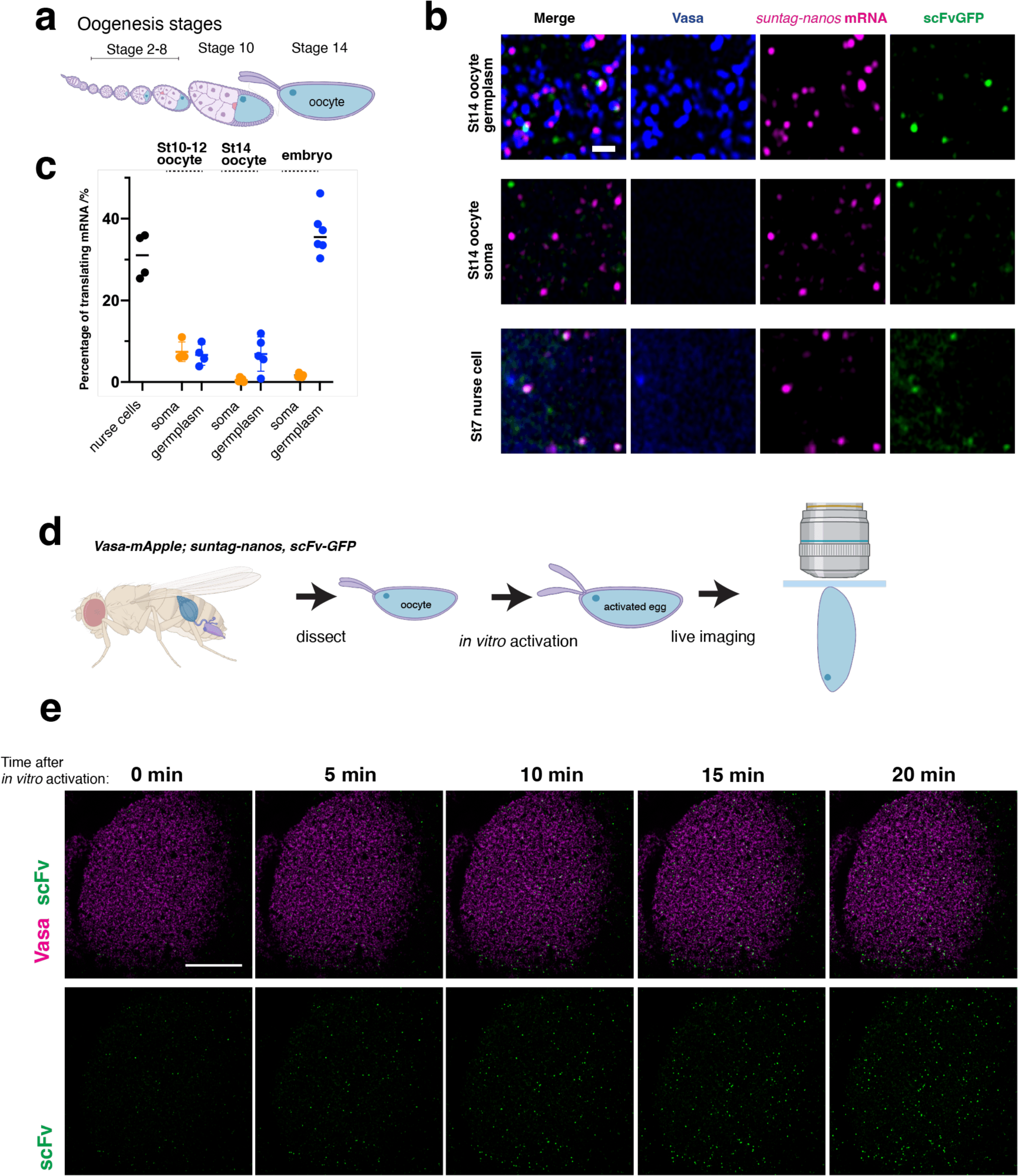
*Suntag-nanos* mRNA translation during oogenesis. **a**, Schematic of Drosophila oogenesis stages. **b**, Representative images of germplasm (top) and soma (middle) in stage 14 oocyte and cytoplasm of stage 7 nurse cells (bottom) expressing *suntag-nanos* and scFv-GFP. Blue, Vasa; magenta, suntag smFISH; green, scFv-GFP. Scale bar 1 µm. **c**, Translating fraction of *suntag-nanos* mRNA in stage 4-10 nurse cells, soma, and germplasm of stage 10-12 (developing) oocytes, stage 14 (mature) oocytes, and stage 1-2 embryos. **d**, Protocol of in vitro activation of oocytes and live imaging. Mature oocytes are dissected from Vasa-mApple/+; *suntag-nanos*, scFv-GFP/+ flies and activated with 30% Robb’s buffer (see method for details). Activated eggs are mounted onto a coverslip and imaged by confocal microscopy. **e**, Representative time-lapse images of the germplasm of an activated egg with an increasing number of polysome (green foci). Germplasm is marked by Vasa-mApple (magenta) and SunTag is detected by endogenous scFv-GFP (green). The top shows the merged image, and the bottom shows scFv-GFP channel only. Scale bar 20 µm. Schematics in **a** and **d** were generated with BioRender (https://www.biorender.com/)

**Extended Data Fig. 5.**
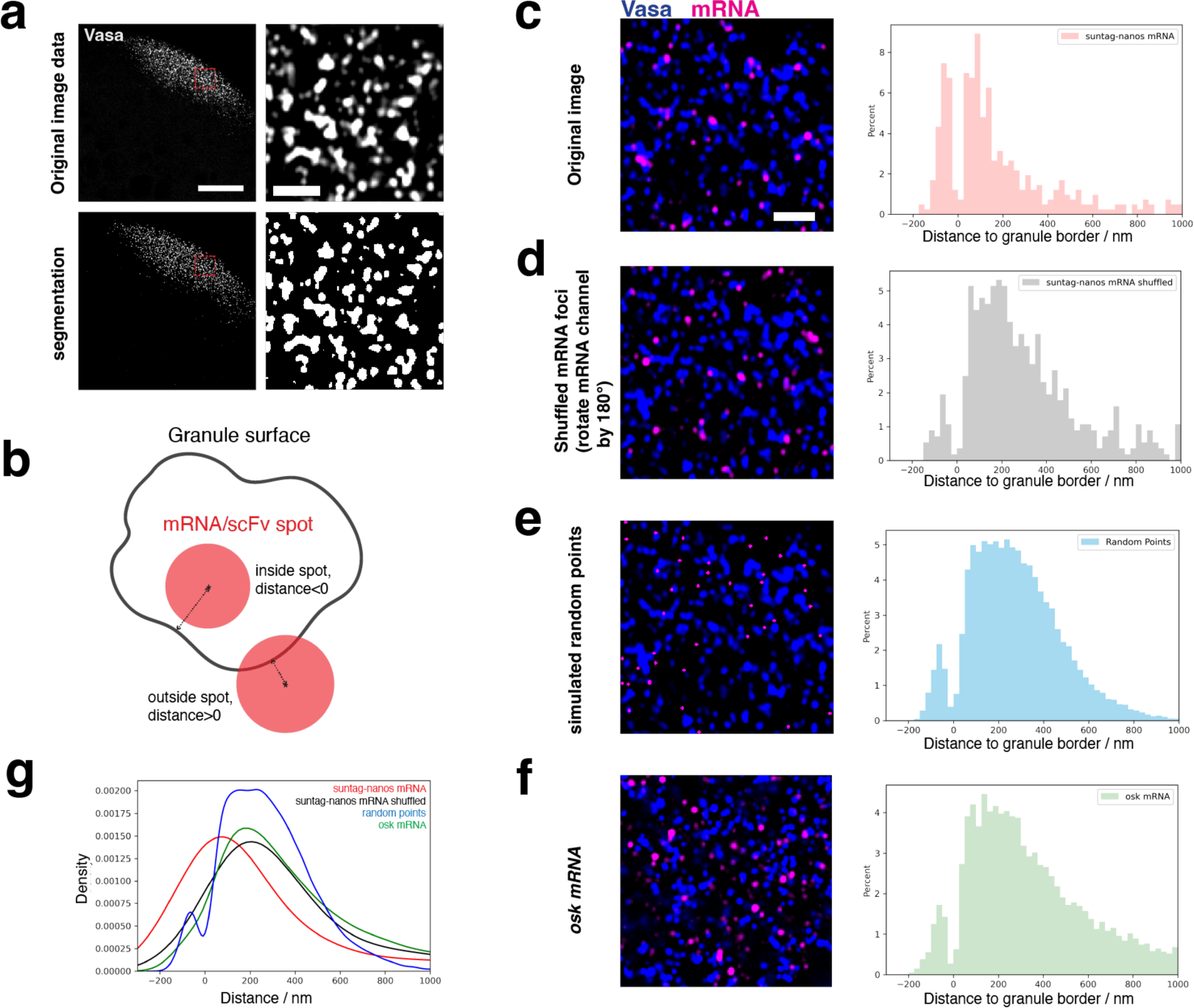
Granule segmentation and distance measurement. **a**, Images of germ granules are segmented with the Ilastik program. The top shows the original grayscale images of Vasa-mApple at the posterior pole of an embryo (left) and zoomed image of the outline region in the germplasm (right). The bottom shows the black-and-white binary images of segmented germ granules (granules in white). Scale bar 20 µm (left), and 2 µm (right). **b**, Schematic of the distance measurement program. The granule surface is defined after segmentation by Ilastik. The coordinates of mRNA smFISH or scFv-GFP/anti-GCN4 spots are determined by FISH-Quant and used to measure to distance to the closest granule surface. The schematic is drawn in 2D but the actual data and measurement are in 3D (see methods). **c**-**f**, Control and validation experiments of distance measurement. (Left) representative images of germplasm with germ granules marked by Vasa-mApple (blue). Magenta: **c** suntag mRNA smFISH; **d** is the same image as **c** with mRNA channel rotated by 180° to shuffle the mRNA distribution; **e** has the same Vasa channel image as **c** with simulated points randomly distributed within the image; **f** smFISH of osk mRNA. Scale bar 2 µm. The distributions of mRNA foci or simulated points are plotted in the relative frequency histograms on the right. The x-axis refers to the distance of foci centroids to the border of the closest granule; the zero marks granule border; a negative value denotes being inside a granule and positive denotes outside. The two bins around 0 have abnormally low counts in all experiments, likely an artifact caused by the design of the distance measuring program. **g** The distributions of mRNA foci or simulated points in (**c**-**f**) are plotted together as a kernel density estimate (KDE) plot. The distributions of shuffled mRNA, random points, and osk mRNA show a shift away from the surface of germ granules when compared to *suntag-nanos* mRNA.

**Extended Data Fig. 6.**
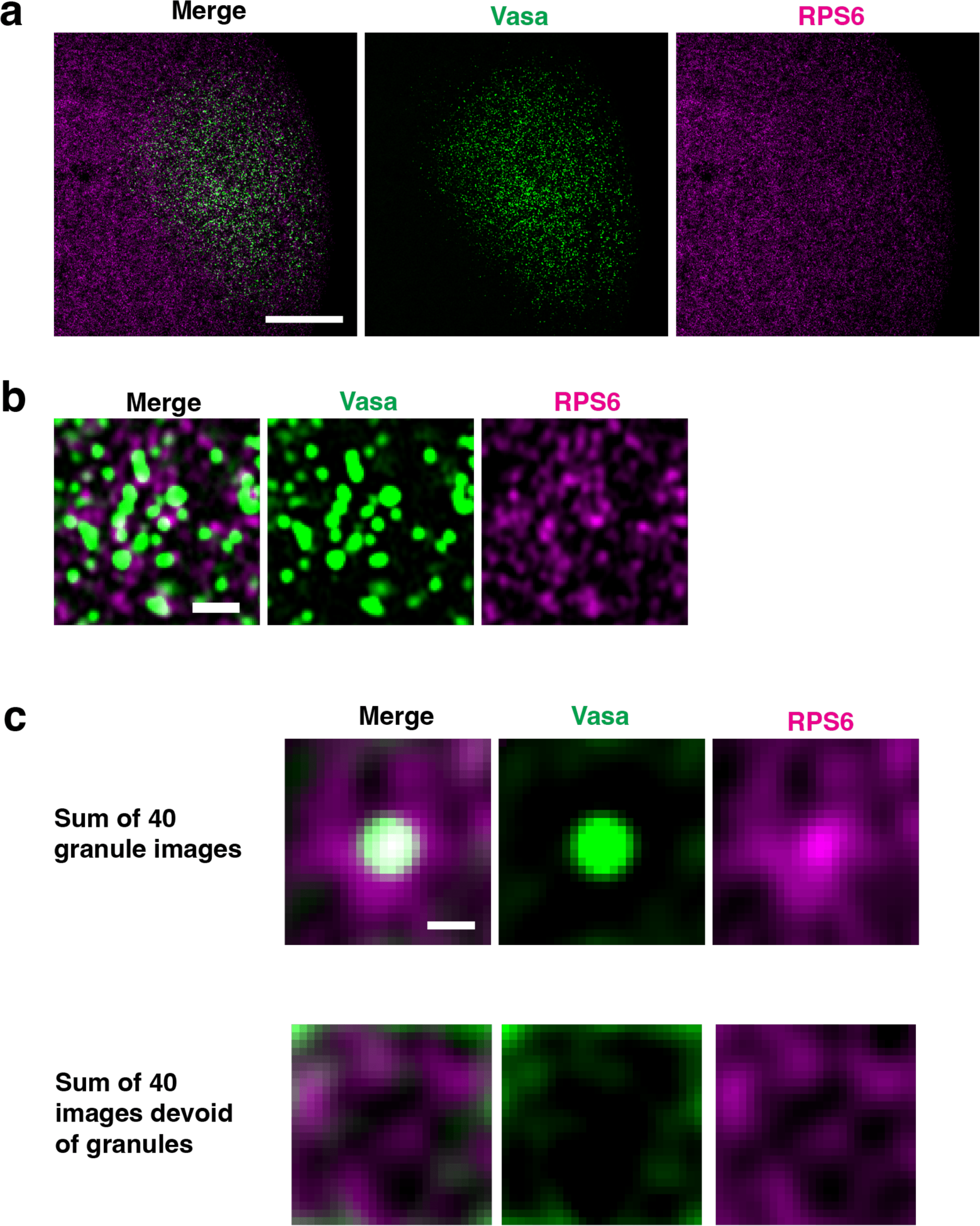
Distribution of ribosomes in germplasm. **a**-**c**, Images of germplasm with germ granules marked by VasaGFP (green) and RPS6 (Ribosomal Protein S6) stained by anti-RPS6 (magenta). (A) shows the posterior pole of the embryo. Scale bar 20 µm. **b**, Zoomed image in germplasm showing the distribution of RPS6. Scale bar 1µm. **c**, Z-stacks of 40 images (26 pixels x 26 pixels) of germplasm with germ granules at the center or without germ granules were made and z-projected by summing slices. Scale bar 0.25 µm.

**Extended Data Fig. 7.**
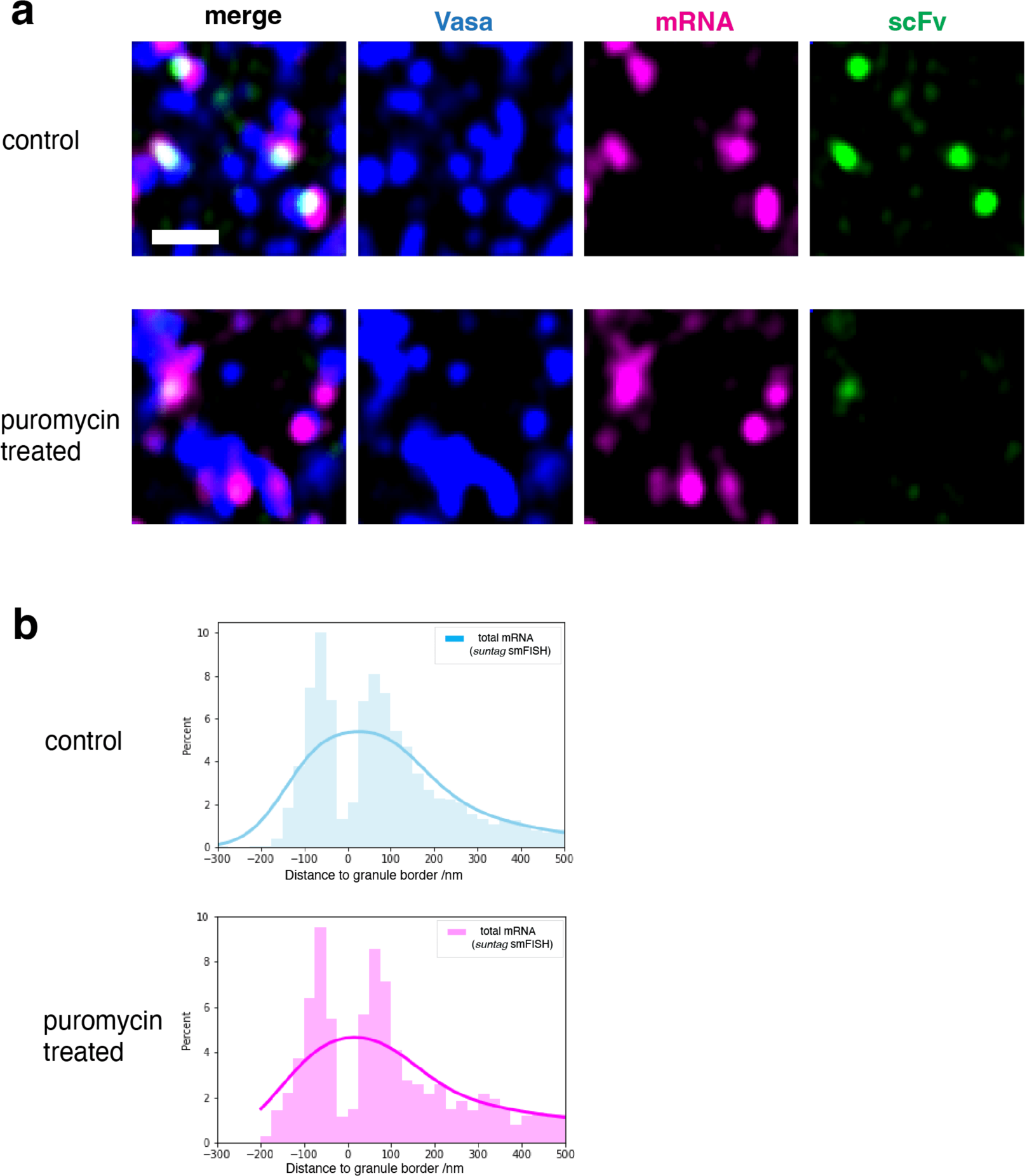
Distribution of suntag CDS is not affected by puromycin treatment. **a**, Distribution of *suntag-nanos* mRNA in germplasm after injecting 20 mM HEPES buffer (control, top) or 10mg/ml puromycin (bottom) and 30 min aging. Blue, Vasa; magenta, *suntag* smFISH; green, scFv-GFP. Scale bar 1 µm. **b**, Distributions of total mRNA detected by suntag smFISH in germplasm of HEPES-injected embryos and puromycin-injected embryos were plotted in relative frequency histograms with KDE curves. Spots from three embryos of each condition were mapped and plotted.

**Extended Data Fig. 8.**
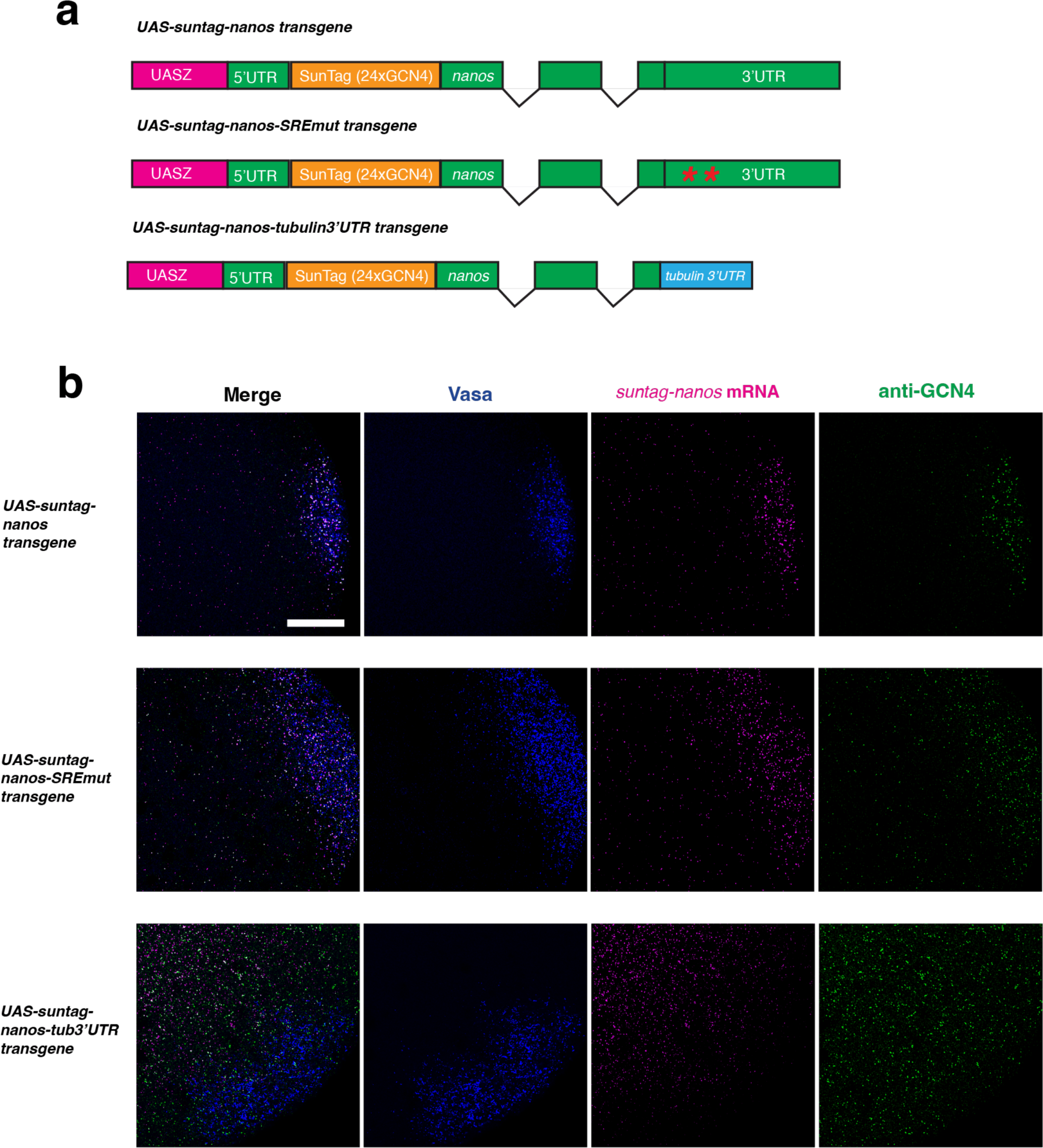
Translation regulation of *suntag-nanos* is mediated by *nanos* 3’UTR. **a**, Schematics of transgenic constructs of *UAS-suntag-nanos*, *UAS-suntag-nanos-SREmut*, and *UAS-suntag-nanos-tubulin3’UTR*. The red asterisks in *nanos* 3’UTR represent the two SREs mutated in the construct. **b**, Representative images of embryos expressing *suntag-nanos* (top), *suntag-nanos-SREmut* (middle), and *suntag-nanos-tubulin3’UTR* (bottom). Note that the translation activities in the soma of embryos expressing *suntag-nanos-SREmut* and *suntag-nanos-tubulin3’UTR* are higher than *suntag-nanos*. Blue, Vasa; magenta, suntag smFISH; green, anti-GCN4. Scale bar 20 µm.

**Extended Data Fig. 9.**
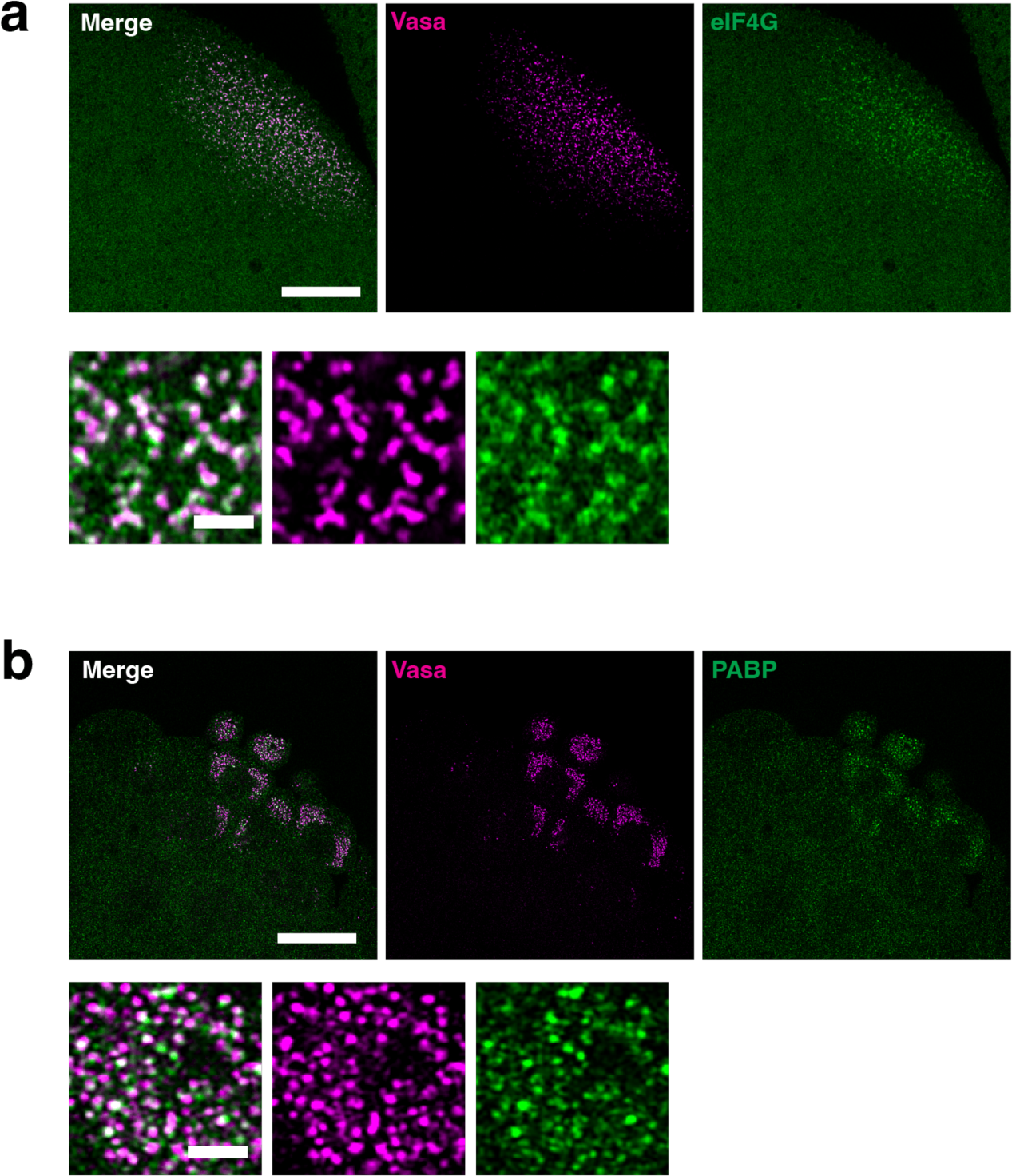
Localization of translation factors to germ granules. **a**, (Top) the posterior of an embryo expressing Vasa-mCherry and eIF4G-YFP, and zoomed images of germplasm (bottom), showing the enrichment of eIF4G (green) to germ granules (Vasa-mCherry, magenta). **b**, (Top) the posterior of an embryo expressing Vasa-mCherry and PABP-YFP, and zoomed images of germplasm (bottom), showing the association of PABP puncta (green) with germ granules (Vasa-mCherry, magenta). Scale bar: top 20 µm, bottom 2 µm.

**Extended Data Fig. 10.**
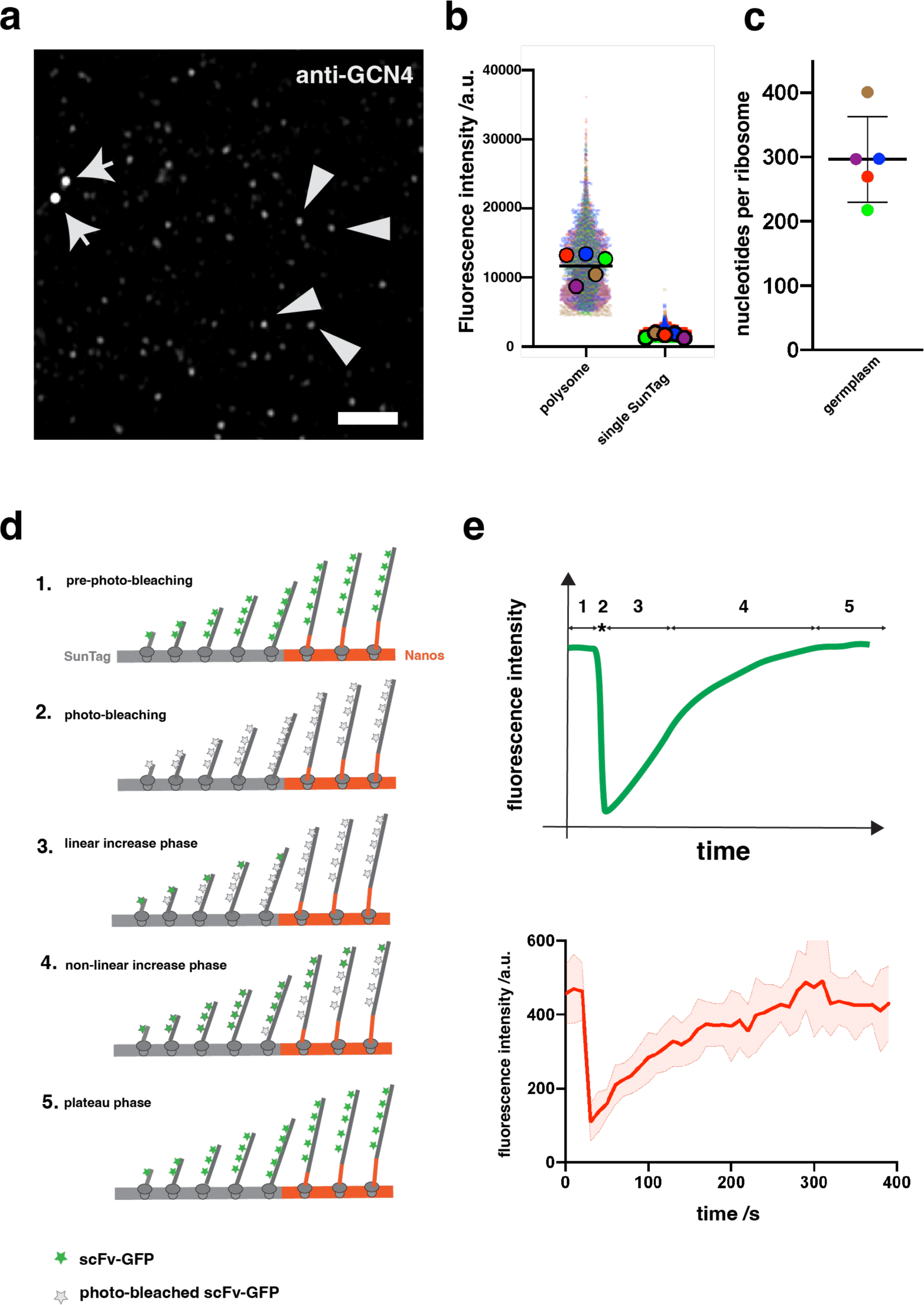
Quantification of the intensity of polysomes, ribosome occupancy, and translation elongation. **a**, Germplasm of an embryo expressing *suntag-nanos* flies with SunTag detected by anti-GCN4. The polysomes (arrows) have stronger fluorescence intensities and are co-localized with the mRNA signal (not shown). Individual synthesized SunTag-Nanos proteins (examples pointed out by arrowheads) have lower intensities and are not co-localized with mRNA. Scale bar 2 µm. **b**, Fluorescence intensities of polysomes and single SunTag-Nanos protein, extracted from FISH-Quant analysis (see methods). Data from five embryos, represented by different colors, are plotted as a super-plot. **c**, Calculated ribosome occupancy on *suntag-nanos* mRNA using data from **b**. **d**, Theoretical process of fluorescence recovery after photo-bleaching (FRAP). Before photo-bleaching, *suntag-nanos* mRNA is translated at a steady state with SunTag bound by fluorescent scFv-GFP (phase 1). Photo-bleaching diminishes the fluorescence of bound scFv-GFP (phase 2). Newly synthesized SunTag epitopes after photo-bleaching bind fluorescent scFv-GFP, causing fluorescence recovery of the polysome. Assuming a constant elongation rate, the initial phase of recovery is linear (phase 3). When the peptide that contains the first SunTag synthesized post-bleaching leaves polysome, which counteracts the increase of newly synthesized SunTag, the increase of signal starts to slow down (phase 4). When the first ribosome loaded after photo-bleaching finishes the translation, the signal reaches a plateau (phase 5) with the same intensity as before photo-bleaching because all the SunTags are bound by fluorescent scFv-GFP again. **e**, A hypothetical FRAP curve (top) based on the theoretical FRAP process, and the FRAP experimental data (bottom, same as Figure 3e), which shows a similar curve as the theoretical curve.

**Extended Data Fig. 11.**
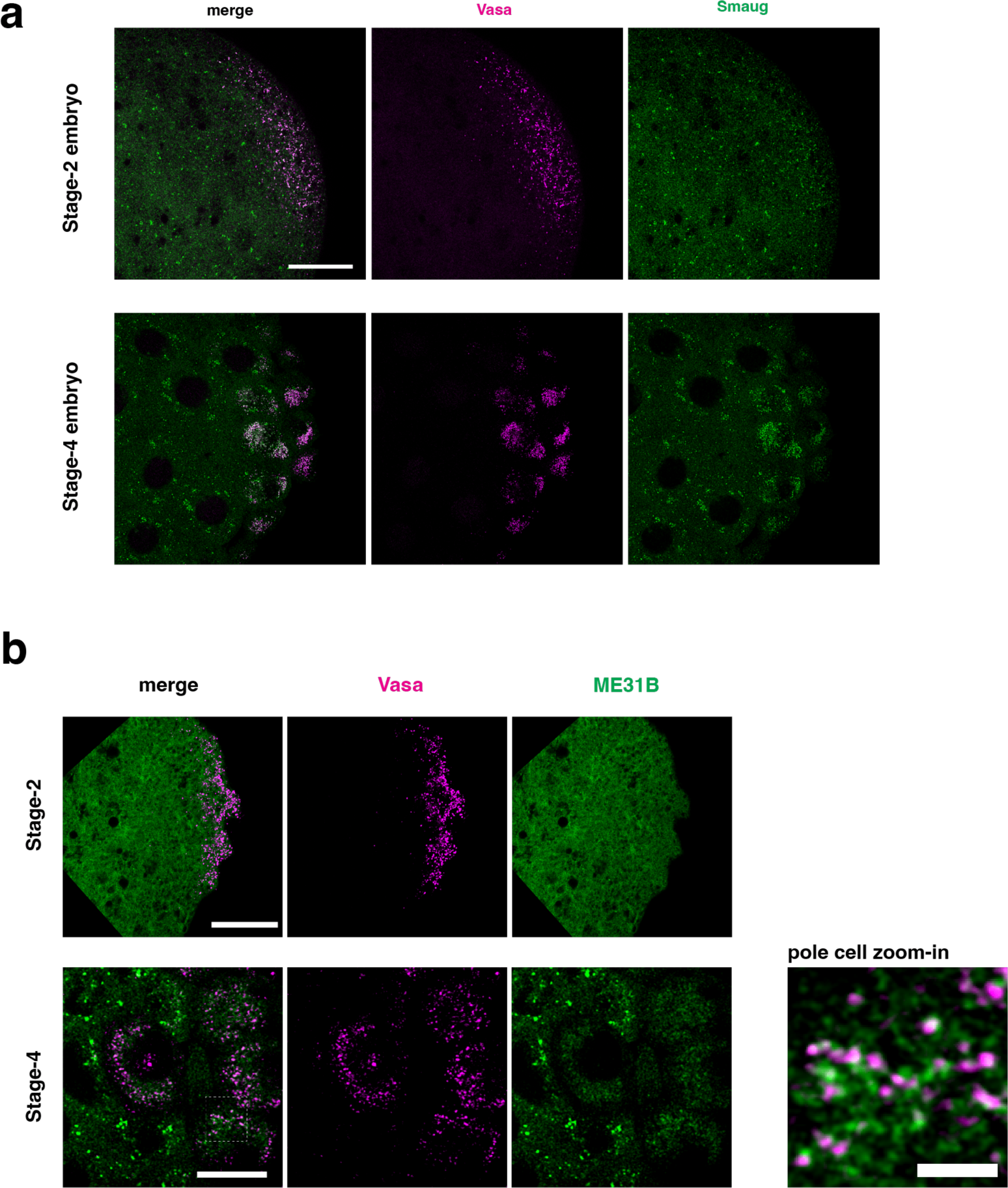
Distribution of Smaug and ME31B. **a**, Stage-2 (top) and stage-4 (bottom) embryos expressing Vasa-mApple and Smaug-GFP, showing the morphology and distribution of Smaug (green) in soma and germplasm. In the soma, Smaug forms heterogeneous puncta. In germplasm, Smaug is enriched in germ granules (magenta). Scale bar 20 µm. **b**, Stage-2 (top) and stage-4 (bottom) embryos expressing Vasa-mApple and ME31B-GFP, showing the distribution of ME31B (green). At stage 2, ME31B is homogeneously distributed throughout the embryo. At stage 4 and later, ME31B forms large and heterogeneous clusters in the soma and forms small clusters associated with germ granules (magenta) in pole cells, as shown in the zoomed image of the outlined area. Scale bar: top 20 µm, bottom 10 µm, zoomed-in image 5 µm.

**Extended Data Fig. 12.**
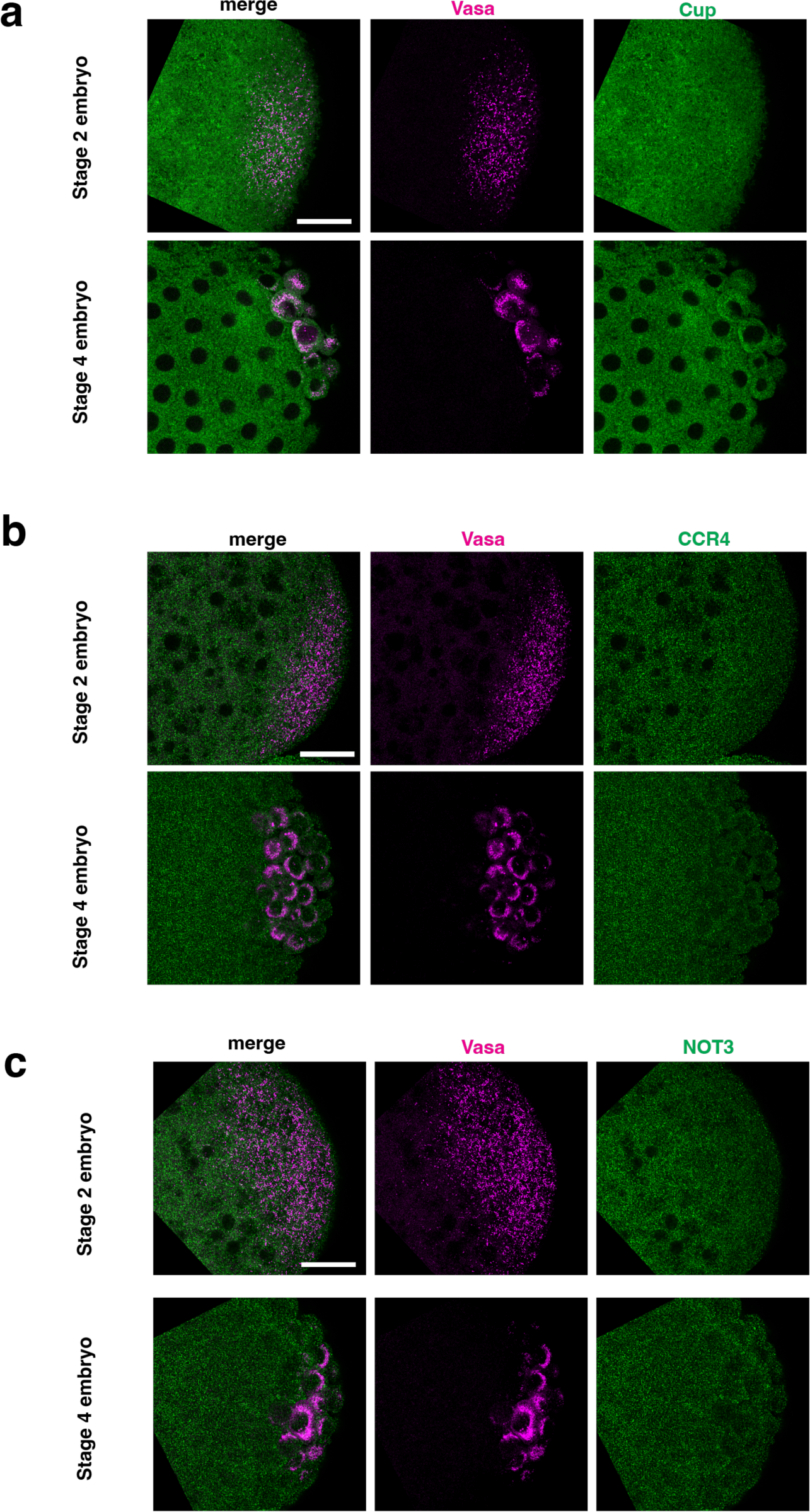
Distribution of Cup, CCR4, and NOT3. **a**, Embryos expressing Cup-YFP (green) and Vasa-mCherry (magenta). **b**, Embryos expressing Vasa-mApple (magenta) stained with anti-CCR4 antibody (green). **c**, Embryos expressing Vasa-mApple (magenta) stained with anti-NOT3 antibody (green). Stage-2 embryos are shown on the top and stage-4 embryos are shown at the bottom. Scale bar 20 µm.

**Extended Data Fig. 13.**
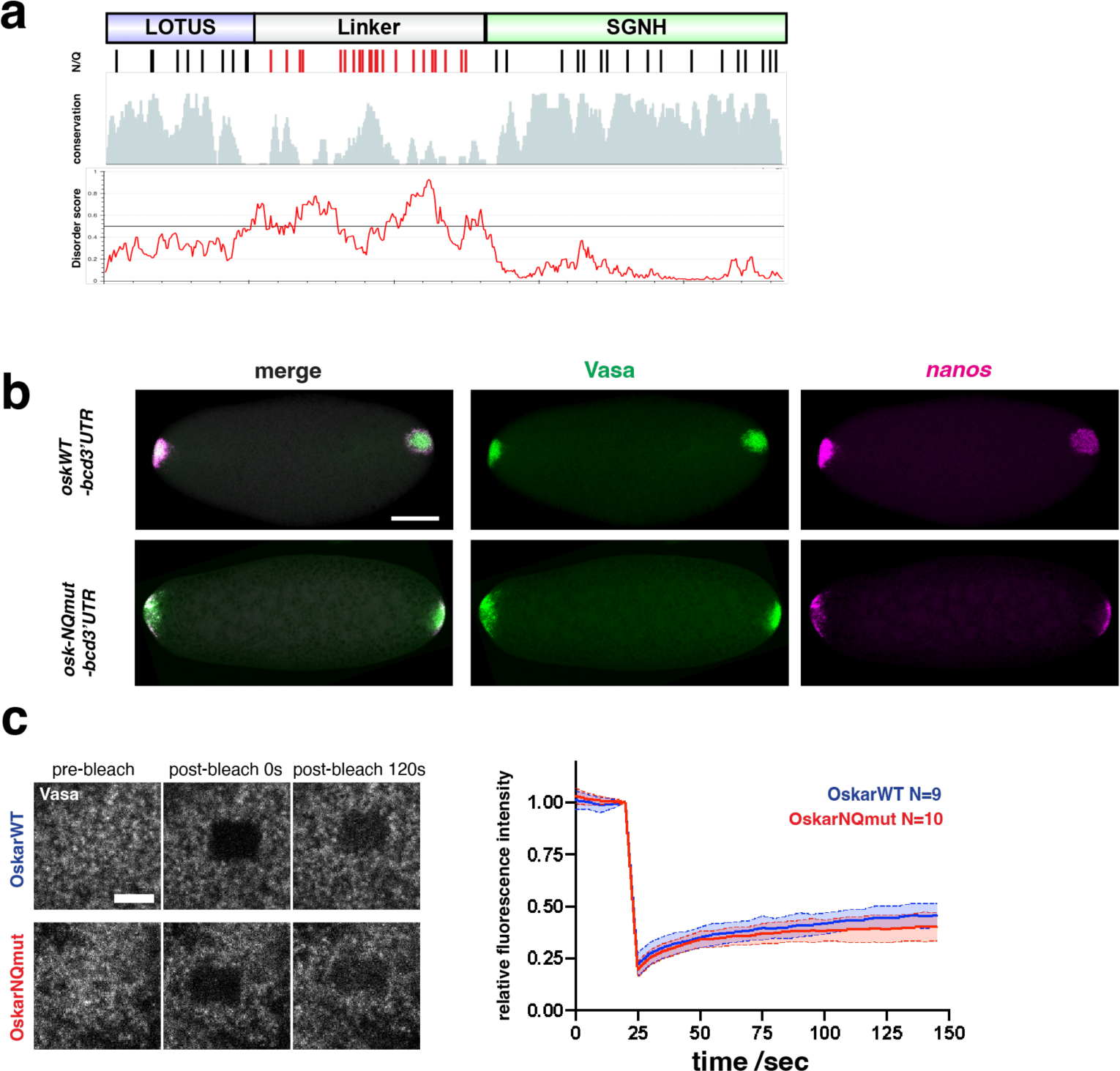
Characterization of Oskar-NQmut. **a**, Sequence features of short Oskar protein. (Top to bottom) The first track shows the domain structure of Oskar. The second track shows the distribution of Asparagine (N) and Glutamine (Q) residues in the Oskar of Drosophila melanogaster. The third track shows the sequence conservation of Oskar proteins of 11 Drosophila species. The fourth track shows disorder prediction of the Oskar sequence using IUPred2A online tool. **b**, Embryos expressing *oskWT-bcd3’UTR* (top) or *osk-NQmut-bcd3’UTR* (bottom). Germplasm is marked by Vasa (green) and *nanos* mRNA is stained by smFISH (magenta). Scale bar 100 µm. **c**, FRAP of Vasa-mApple in anterior germplasm of embryos expressing *oskWT-bcd3’UTR* (top) or *osk-NQmut-bcd3’UTR* (bottom). Fluorescence intensity over time (WT: blue; NQmut: red) is plotted on the right. Scale bar 5 µm.

**Table S1.**
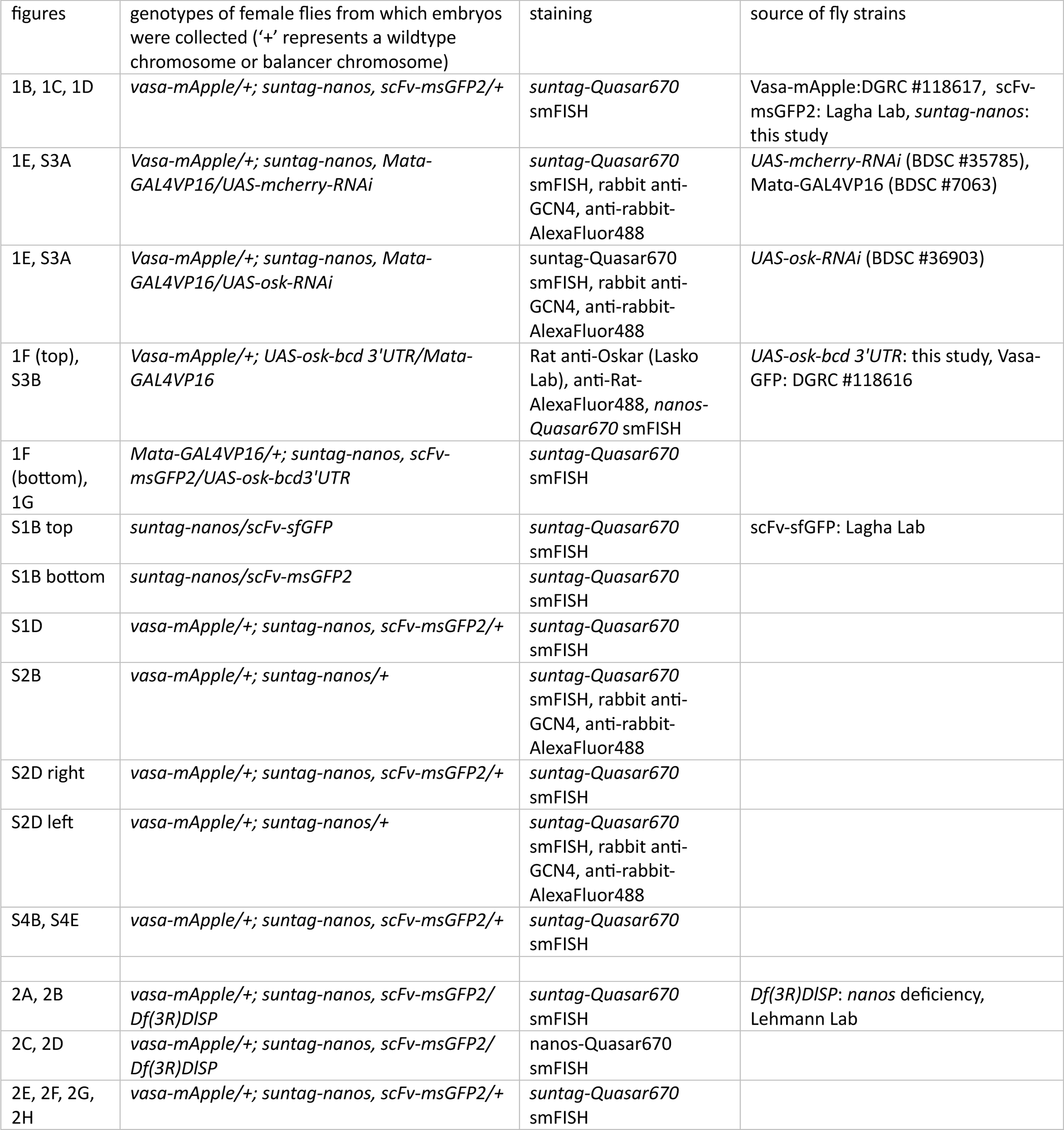
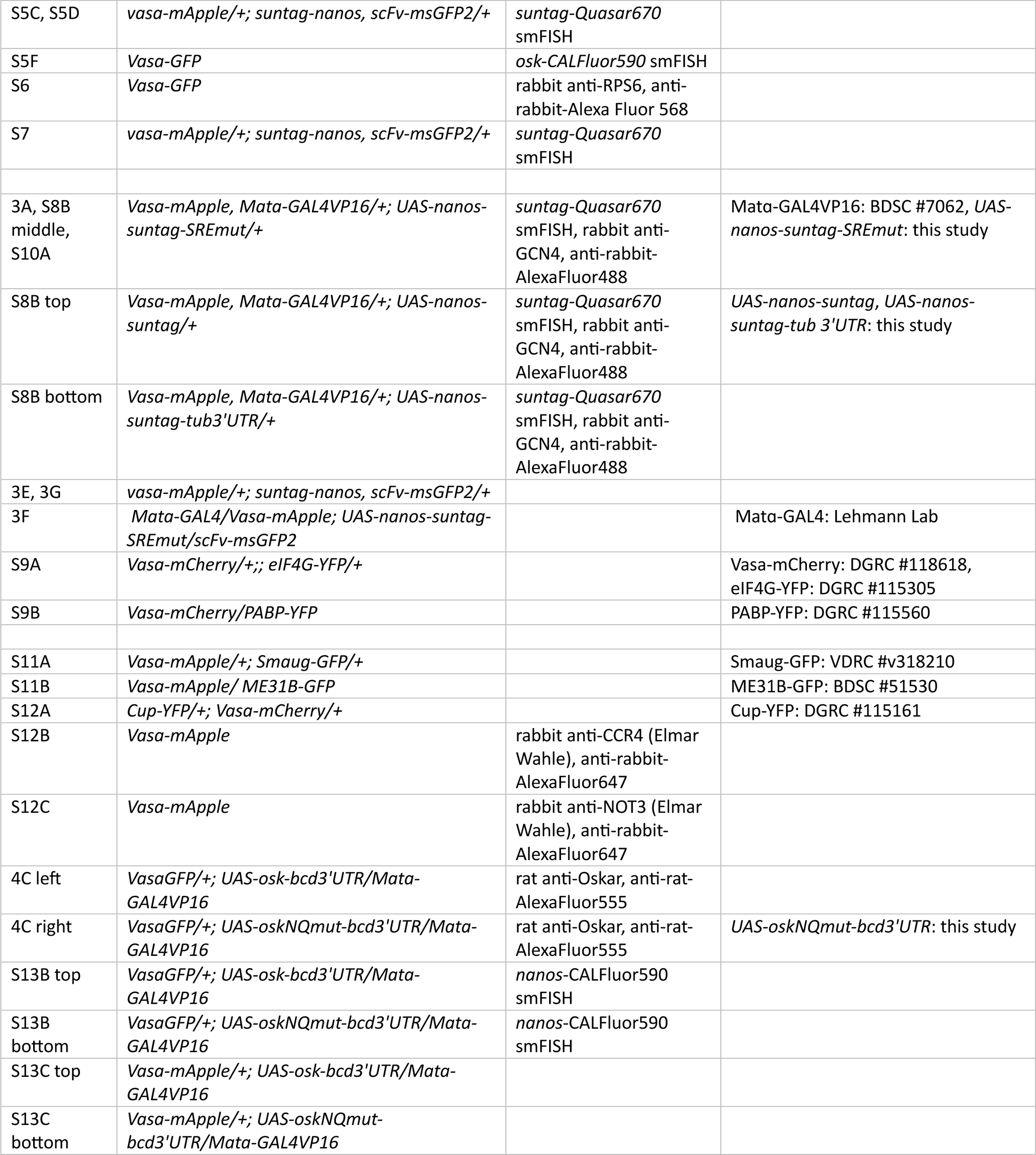
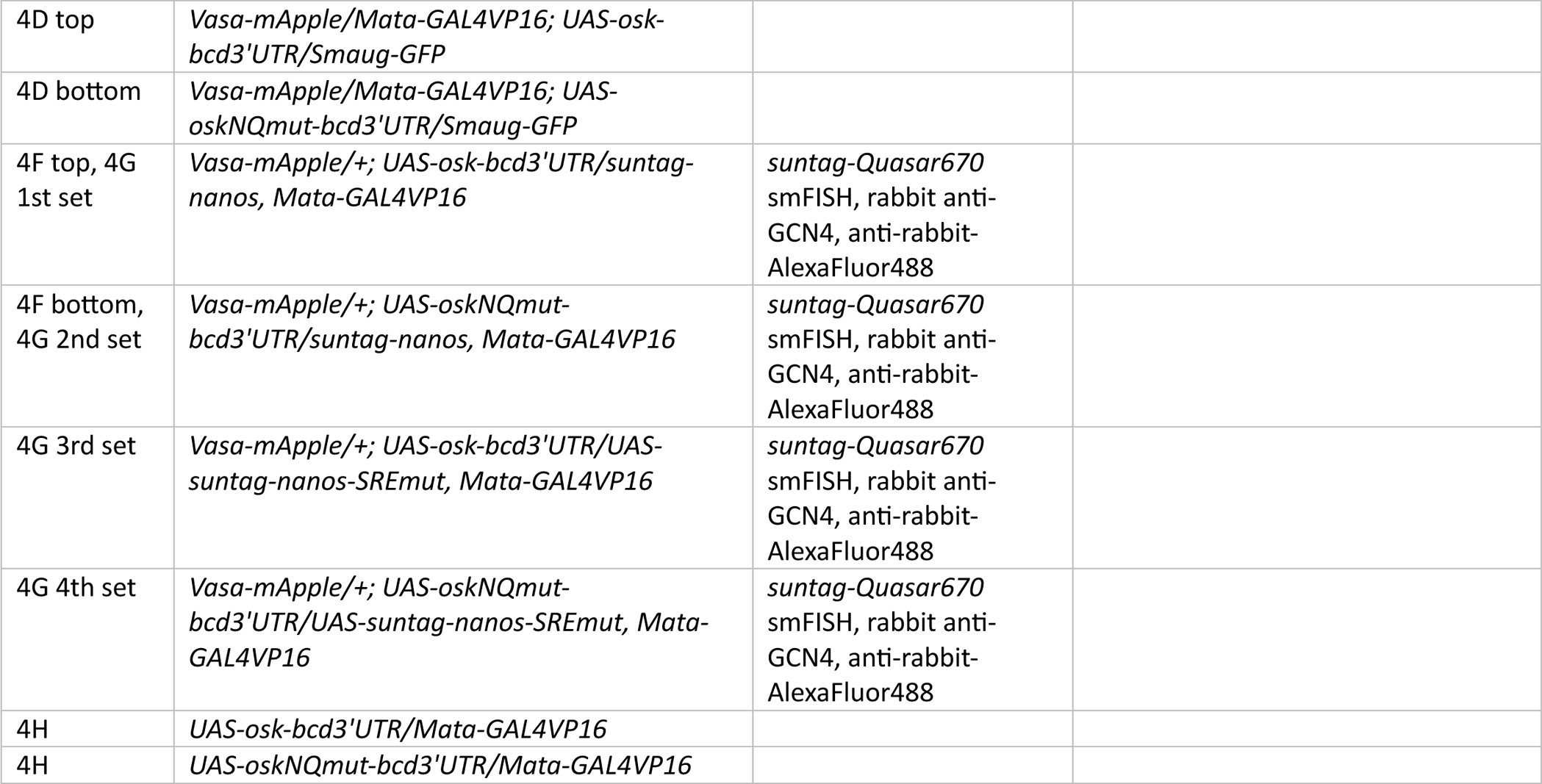
Genotypes of the experimental flies. The methods and reagents used in staining.

| figures | genotypes of female flies from which embryos were collected ('+' represents a wildtype chromosome or balancer chromosome) | staining | source of fly strains |
| --- | --- | --- | --- |
| 1B, 1C, 1D | *vasa-mApple/+; suntag-nanos, scFv-msGFP2/+* | *suntag-Quasar670* smFISH | Vasa-mApple: DGRC #118617, scFv-msGFP2: Lagha Lab, *suntag-nanos*: this study |
| 1E, S3A | *Vasa-mApple/+; suntag-nanos, Mata-GAL4VP16/UAS-mcherry-RNAi* | *suntag-Quasar670* smFISH, rabbit anti-GCN4, anti-rabbit-AlexaFluor488 | *UAS-mcherry-RNAi* (BDSC #35785), Mata-GAL4VP16 (BDSC #7063) |
| 1E, S3A | *Vasa-mApple/+; suntag-nanos, Mata-GAL4VP16/UAS-osk-RNAi* | *suntag-Quasar670* smFISH, rabbit anti-GCN4, anti-rabbit-AlexaFluor488 | *UAS-osk-RNAi* (BDSC #36903) |
| 1F (top), S3B | *Vasa-mApple/+; UAS-osk-bcd 3'UTR/Mata-GAL4VP16* | Rat anti-Oskar (Lasko Lab), anti-Rat-AlexaFluor488, *nanos-Quasar670* smFISH | *UAS-osk-bcd 3'UTR*: this study, Vasa-GFP: DGRC #118616 |
| 1F (bottom), 1G | *Mata-GAL4VP16/+; suntag-nanos, scFv-msGFP2/UAS-osk-bcd3'UTR* | *suntag-Quasar670* smFISH |  |
| S1B top | *suntag-nanos/scFv-sfGFP* | *suntag-Quasar670* smFISH | scFv-sfGFP: Lagha Lab |
| S1B bottom | *suntag-nanos/scFv-msGFP2* | *suntag-Quasar670* smFISH |  |
| S1D | *vasa-mApple/+; suntag-nanos, scFv-msGFP2/+* | *suntag-Quasar670* smFISH |  |
| S2B | *vasa-mApple/+; suntag-nanos/+* | *suntag-Quasar670* smFISH, rabbit anti-GCN4, anti-rabbit-AlexaFluor488 |  |
| S2D right | *vasa-mApple/+; suntag-nanos, scFv-msGFP2/+* | *suntag-Quasar670* smFISH |  |
| S2D left | *vasa-mApple/+; suntag-nanos/+* | *suntag-Quasar670* smFISH, rabbit anti-GCN4, anti-rabbit-AlexaFluor488 |  |
| S4B, S4E | *vasa-mApple/+; suntag-nanos, scFv-msGFP2/+* | *suntag-Quasar670* smFISH |  |
| 2A, 2B | *vasa-mApple/+; suntag-nanos, scFv-msGFP2/Df(3R)DISP* | *suntag-Quasar670* smFISH | *Df(3R)DISP*: *nanos* deficiency, Lehmann Lab |
| 2C, 2D | *vasa-mApple/+; suntag-nanos, scFv-msGFP2/Df(3R)DISP* | *nanos-Quasar670* smFISH |  |
| 2E, 2F, 2G, 2H | *vasa-mApple/+; suntag-nanos, scFv-msGFP2/+* | *suntag-Quasar670* smFISH |  |
| S5C, S5D | *vasa-mApple/+; suntag-nanos, scFv-msGFP2/+* | *suntag-Quasar670*<br>smFISH |  |
| S5F | *Vasa-GFP* | *osk-CALFluor590* smFISH |  |
| S6 | *Vasa-GFP* | rabbit anti-RPS6, anti-rabbit-Alexa Fluor 568 |  |
| S7 | *vasa-mApple/+; suntag-nanos, scFv-msGFP2/+* | *suntag-Quasar670*<br>smFISH |  |
| 3A, S8B middle, S10A | *Vasa-mApple, Mata-GAL4VP16/+; UAS-nanos-suntag-SREmut/+* | *suntag-Quasar670* smFISH, rabbit anti-GCN4, anti-rabbit-AlexaFluor488 | Mata-GAL4VP16: BDSC #7062, *UAS-nanos-suntag-SREmut*: this study |
| S8B top | *Vasa-mApple, Mata-GAL4VP16/+; UAS-nanos-suntag/+* | *suntag-Quasar670* smFISH, rabbit anti-GCN4, anti-rabbit-AlexaFluor488 | *UAS-nanos-suntag, UAS-nanos-suntag-tub 3'UTR*: this study |
| S8B bottom | *Vasa-mApple, Mata-GAL4VP16/+; UAS-nanos-suntag-tub3'UTR/+* | *suntag-Quasar670* smFISH, rabbit anti-GCN4, anti-rabbit-AlexaFluor488 |  |
| 3E, 3G | *vasa-mApple/+; suntag-nanos, scFv-msGFP2/+* |  |  |
| 3F | *Mata-GAL4/Vasa-mApple; UAS-nanos-suntag-SREmut/scFv-msGFP2* |  | Mata-GAL4: Lehmann Lab |
| S9A | *Vasa-mCherry/+;; eIF4G-YFP/+* |  | Vasa-mCherry: DGRC #118618, eIF4G-YFP: DGRC #115305 |
| S9B | *Vasa-mCherry/PABP-YFP* |  | PABP-YFP: DGRC #115560 |
| S11A | *Vasa-mApple/+; Smaug-GFP/+* |  | Smaug-GFP: VDRC #v318210 |
| S11B | *Vasa-mApple/ ME31B-GFP* |  | ME31B-GFP: BDSC #51530 |
| S12A | *Cup-YFP/+; Vasa-mCherry/+* |  | Cup-YFP: DGRC #115161 |
| S12B | *Vasa-mApple* | rabbit anti-CCR4 (Elmar Wahle), anti-rabbit-AlexaFluor647 |  |
| S12C | *Vasa-mApple* | rabbit anti-NOT3 (Elmar Wahle), anti-rabbit-AlexaFluor647 |  |
| 4C left | *VasaGFP/+; UAS-osk-bcd3'UTR/Mata-GAL4VP16* | rat anti-Oskar, anti-rat-AlexaFluor555 |  |
| 4C right | *VasaGFP/+; UAS-oskNQmut-bcd3'UTR/Mata-GAL4VP16* | rat anti-Oskar, anti-rat-AlexaFluor555 | *UAS-oskNQmut-bcd3'UTR*: this study |
| S13B top | *VasaGFP/+; UAS-osk-bcd3'UTR/Mata-GAL4VP16* | *nanos-CALFluor590* smFISH |  |
| S13B bottom | *VasaGFP/+; UAS-oskNQmut-bcd3'UTR/Mata-GAL4VP16* | *nanos-CALFluor590* smFISH |  |
| S13C top | *Vasa-mApple/+; UAS-osk-bcd3'UTR/Mata-GAL4VP16* |  |  |
| S13C bottom | *Vasa-mApple/+; UAS-oskNQmut-bcd3'UTR/Mata-GAL4VP16* |  |  |
| 4D top | *Vasa-mApple/Mata-GAL4VP16; UAS-osk-bcd3'UTR/Smaug-GFP* |  |  |
| 4D bottom | *Vasa-mApple/Mata-GAL4VP16; UAS-oskNQmut-bcd3'UTR/Smaug-GFP* |  |  |
| 4F top, 4G 1st set | *Vasa-mApple/+; UAS-osk-bcd3'UTR/suntag-nanos, Mata-GAL4VP16* | *suntag-Quasar670*<br>smFISH, rabbit anti-GCN4, anti-rabbit-AlexaFluor488 |  |
| 4F bottom, 4G 2nd set | *Vasa-mApple/+; UAS-oskNQmut-bcd3'UTR/suntag-nanos, Mata-GAL4VP16* | *suntag-Quasar670*<br>smFISH, rabbit anti-GCN4, anti-rabbit-AlexaFluor488 |  |
| 4G 3rd set | *Vasa-mApple/+; UAS-osk-bcd3'UTR/UAS-suntag-nanos-SREmut, Mata-GAL4VP16* | *suntag-Quasar670*<br>smFISH, rabbit anti-GCN4, anti-rabbit-AlexaFluor488 |  |
| 4G 4th set | *Vasa-mApple/+; UAS-oskNQmut-bcd3'UTR/UAS-suntag-nanos-SREmut, Mata-GAL4VP16* | *suntag-Quasar670*<br>smFISH, rabbit anti-GCN4, anti-rabbit-AlexaFluor488 |  |
| 4H | *UAS-osk-bcd3'UTR/Mata-GAL4VP16* |  |  |
| 4H | *UAS-oskNQmut-bcd3'UTR/Mata-GAL4VP16* |  |  |

**Table S2.**
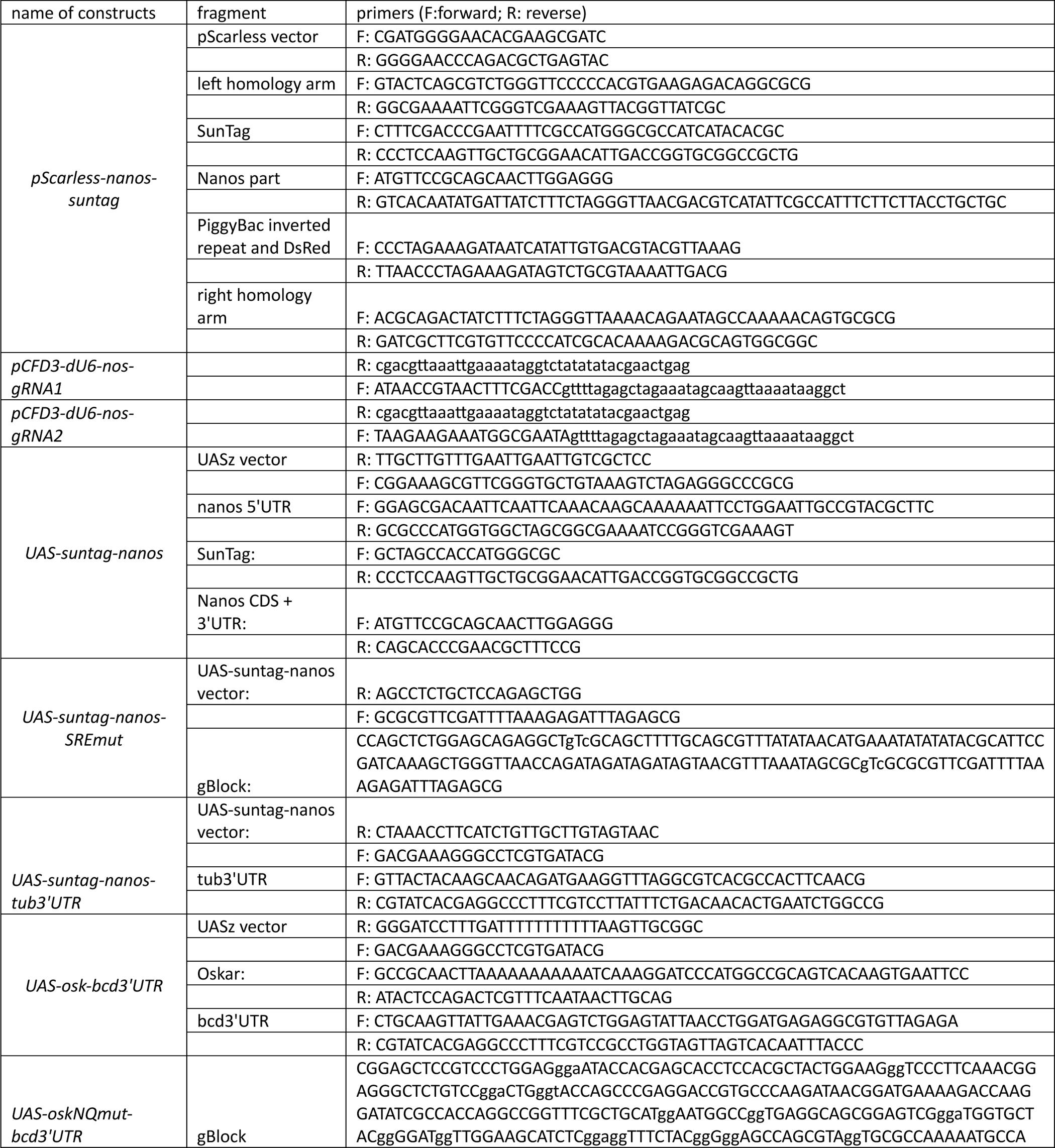
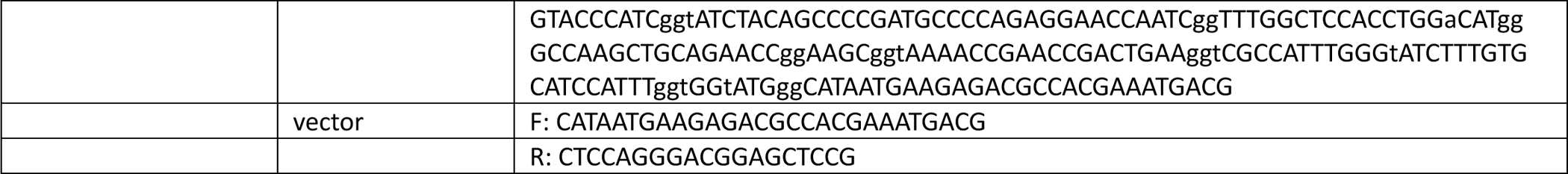
Primers and gBlocks used in cloning.

**Table S3.**
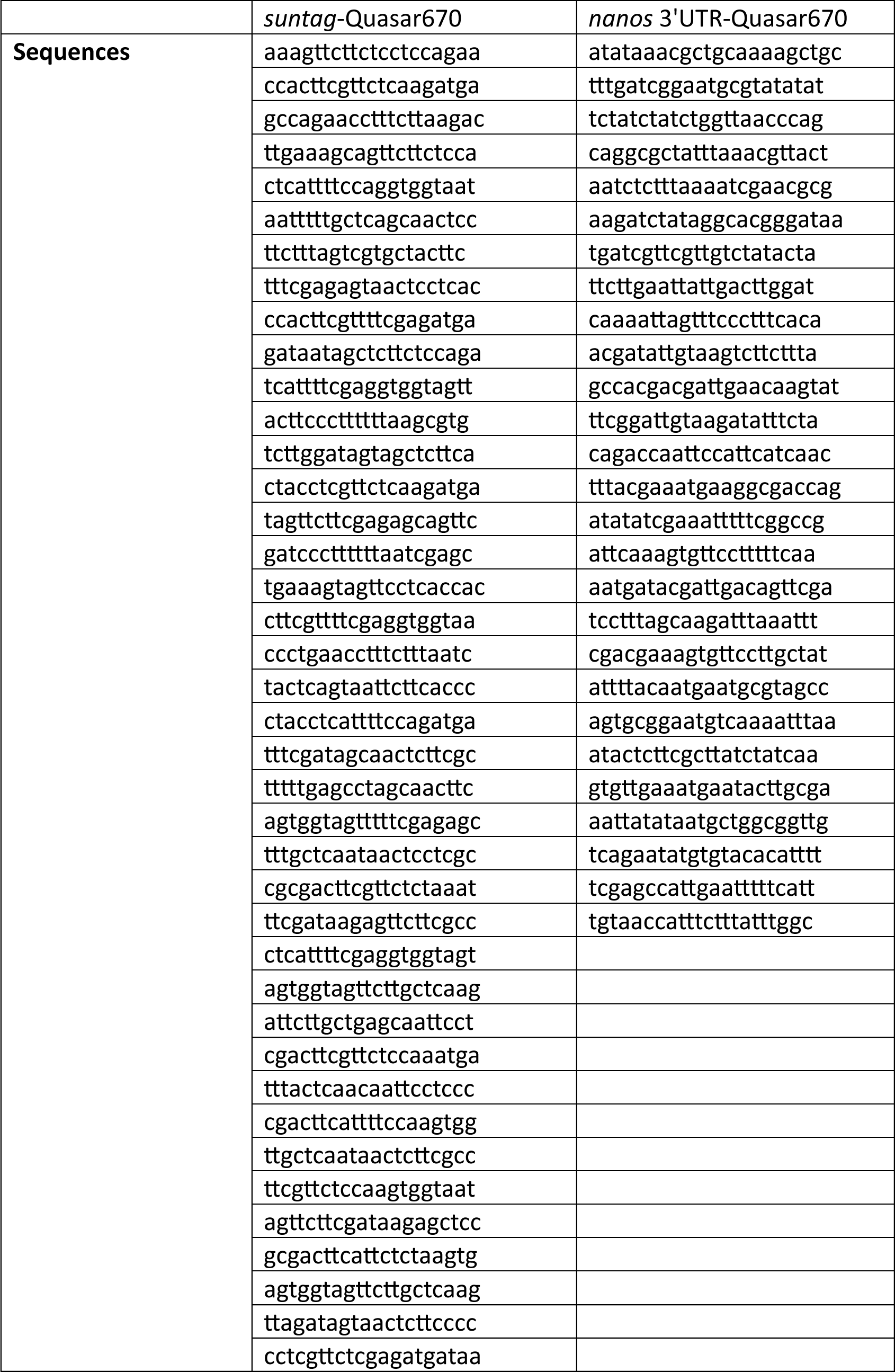
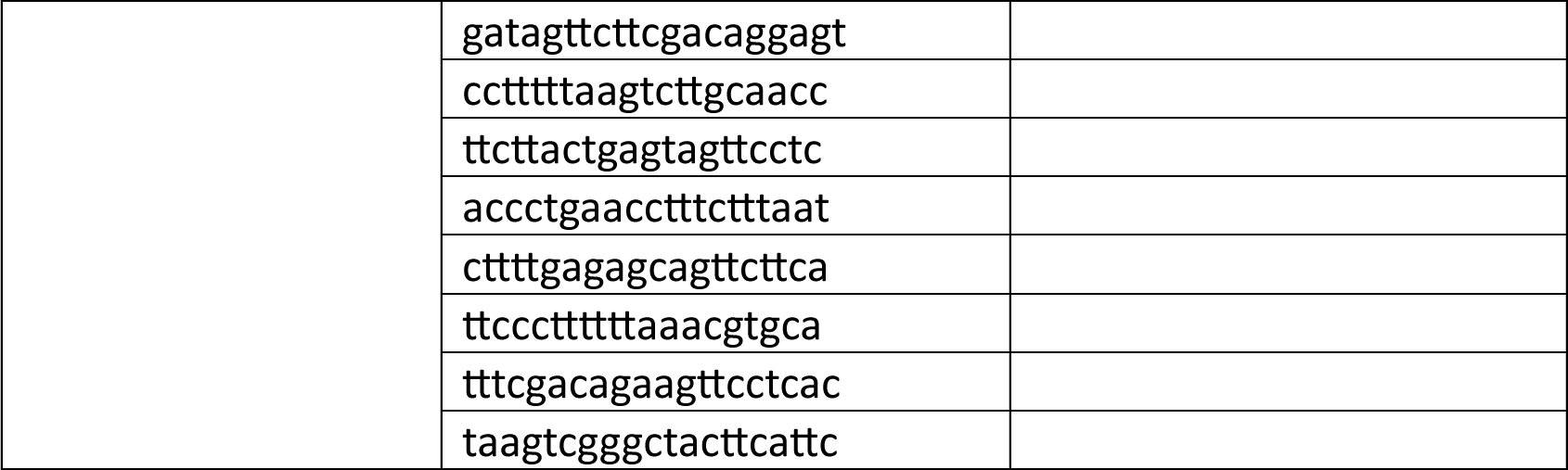
Sequence of smFISH probes.

#### Caption of supplementary movies

**Movie S1.**

Increasing translation in the germ plasm of an *in vitro* activated egg. Stage 14 oocyte was activated in vitro for 30 min before being mounted with posterior stuck on the coverslip and imaged. The timestamp indicates the time after imaging. Germplasm is marked by Vasa-mApple (magenta, middle), SunTag is labeled by scFv-GFP (green, right). Merged image is on the left.

**Movie S2.**

The dynamics of *suntag-nanos* mRNA translation spots over five minutes. SunTag is labeled by scFv-GFP. Note that, despite of constant movement, some of the translation spots (polysomes) stayed within the field of view throughout the imaging process, allowing spot-tracking and intensity measurement over time. Scale bar 1µm.

**Movie S3.**

Example FRAP movie of translation spots in germplasm. SunTag is labeled by scFv-GFP. Three translation spots (arrows) were bleached at 40 sec. The fluorescence of bleached translation spots recovered over time. Scale bar 1µm.

**Movie S4.**

Example FRAP movie of a translation spot in soma. SunTag is labeled by scFv-GFP. One translation spot (arrow) was bleached at 40 sec. The fluorescence of the bleached translation spot recovered over time. Scale bar 1µm.

